# A novel antagonist of the CCL5/CCR5 axis suppresses the tumor growth and metastasis of triple-negative breast cancer by CCR5-YAP1 regulation

**DOI:** 10.1101/2023.11.15.567291

**Authors:** Ling Chen, Guiying Xu, Xiaoxu Song, Lianbo Zhang, Chuyu Chen, Gang Xiang, Shuxuan Wang, Zijian Zhang, Fang Wu, Xuanming Yang, Lei Zhang, Xiaojing Ma, Jing Yu

## Abstract

Triple-negative breast cancer (TNBC) is the most aggressive subtype of breast cancer (BC) with a high mortality rate, and few effective therapeutic strategies are available. CCL5/CCR5 is an appealing immunotherapeutic target for TNBC. However, its signaling mechanism is poorly understood and its direct antagonists have not been reported. Here, we developed a high-throughput screening (HTS) assay for discovering its antagonists. Verteporfin was identified as a more selective and potent antagonist than the known CCR5 antagonist maraviroc. Without photodynamic therapy, verteporfin demonstrated significant inhibition on TNBC tumor growth through immune regulation, remarkable suppression of lung metastasis by cell-intrinsic mechanism, and a significant extension of overall survival *in vivo*. Mechanistically, CCR5 was found to be essential for expression of the key hippo effector YAP1. It promoted *YAP1* transcription via HIF-1α and exerted further control over the migration of CD8^+^ T, NK, and MDSC immune cells through chemokines CXCL16 and CXCL8 which were identified from RNA-seq. Moreover, the CCR5-YAP1 axis played a vital role in promoting metastasis by modulating β-catenin and core epithelial-mesenchymal transition transcription factors ZEB1 and ZEB2. It is noteworthy that the regulatory relationship between CCR5 and YAP1 was observed across various BC subtypes, TNBC patients, and showed potential relevance in fifteen additional cancer types. Overall, this study introduced an easy-to-use HTS assay that streamlines the discovery of CCL5/CCR5 axis antagonists. Verteporfin was identified as a specific molecular probe of this axis with great potentials as a therapeutic agent for treating sixteen malignant diseases characterized by heightened CCR5 and YAP1 levels.

## 1. Introduction

Breast cancer has become the most commonly diagnosed and the fifth leading cause of cancer-related deaths worldwide, surpassing lung cancer in incidence [1]. Among breast cancer cases, triple-negative breast cancer (TNBC) represents approximately 10-20% and is currently the most aggressive subtype lacking effective targeted therapies [2].

In recent years, the C-C chemokine ligand 5/C-C chemokine receptor 5 (CCL5/CCR5) axis has emerged as an appealing therapeutic target in various malignant diseases, including solid tumors, hematological cancers, and even coronavirus disease 2019 (COVID-19) [3-11]. This axis plays a critical role in cancer progression by influencing tumor growth, extracellular matrix remodeling, cancer stem cell expansion, and drug resistance [8]. Additionally, the CCL5/CCR5 axis is implicated in diseases beyond cancer, such as diabetes and Alzheimer’s disease [12, 13]. The CCL5/CCR5 axis does not only have an impact on tumor cells, but also modulates immune cells within the tumor microenvironment [14, 15]. However, the available tools for studying the function and mechanisms of the CCL5/CCR5 axis are limited, the current main approaches were relying on *CCL5*-targeted small interfering RNA (siRNA) or anti-CCL5 antibodies, in combination with or without the only FDA-approved CCR5 antagonist, maraviroc [6, 16]. The absence of specific antagonists for the CCL5/CCR5 axis significantly hinders our understanding of its pathological activities and mechanisms across various diseases.

To date, only a few pathways, like the phosphatidylinositol 3-kinase/protein kinase B (PI3K/PKB) pathway, have been reported to be regulated by the CCL5/CCR5 axis in cancers [5, 17]. However, the precise molecular mechanisms and specific antagonists targeting this axis in TNBC tumor growth and metastasis remain poorly understood.

In this study, we have developed a cell-based high-throughput screening (HTS) assay to identify the antagonists of the CCL5/CCR5 axis for TNBC immunotherapy. By employing a drug repositioning approach, HTS, and multiple preclinical mouse models, we have identified verteporfin as not only a molecular probe for the CCL5/CCR5 axis, but also an immunotherapeutic agent for TNBC and other malignant cancers, including colon adenocarcinoma, pancreatic adenocarcinoma, low-grade glioma, and thymoma. Furthermore, we have unraveled the crucial role of CCR5 in regulating YAP1 oncogene, and elucidated the immune-dependent and cell-intrinsic mechanisms underlying the impact of the CCL5/CCR5 axis on TNBC tumor growth and metastasis.

## 2. Materials and methods

### 2.1. Cell culture

4T1 (CRL-2539, RRID: CVCL_0125), BT549 (HTB-122, RRID: CVCL_1092), HEK293T (CRL-3216, RRID: CVCL_0063), and HCT-116 (CCL-247, RRID: CVCL_0291) cell lines were purchased from the American Type Culture Collection (ATCC, USA). MCF-7 (TCHu 74, RRID: CVCL_0031) and MDA-MB-231 (TCHu 227, RRID: CVCL_0062) cell lines were purchased from the Cell Bank of the Chinese Academy of Sciences (China). Cell lines were authenticated by the vendors. All cell lines were routinely tested for mycoplasma contamination and cultured according to standard protocols. In detailed, the HEK293T, MDA-MB-231, BT549, and MCF-7 cells were grown in Gibco^TM^ BASIC DMEM, High Glucose (ThermoFisher Scientific, C11965500BT), HCT-116 cells were grown in Gibco^TM^ McCoy’s 5A Medium (ThermoFisher Scientific, 16600082), and 4T1 cells were cultured in Gibco^TM^ BASIC RPMI 1640 Medium (ThermoFisher Scientific, C11875500BT). All medium were supplemented with 10% fetal bovine serum (ExCell, FSP500) and 100 μg/ml penicillin/streptomycin (ThermoFisher Scientific, 15140122). All cell lines were maintained at 37°C in a humidified atmosphere containing 95% air and 5% CO2 with the medium changed every two days.

### 2.2. Animal models

All mouse studies were conducted in accordance with the protocols approved by the Institutional Animal Care and Use Committee of Shanghai Jiao Tong university (SJTU, Shanghai, China) (Approval number 202003021). The investigators were blinded to group allocation and data collection for *in vivo* experiments. Mice were maintained in specific pathogen-free conditions with controlled temperature/humidity (22°C/55%) environment on a 12 h light-dark cycle and with standard diet and water ad libitum. The general condition of mice was monitored using their fitness and weight controls. Mice were randomly assigned to experimental groups.

#### 2.2.1. TNBC tumor growth model in an immune competent mouse

2 × 10^5^ 4T1 cells were resuspended in 100 μl PBS and injected into the fourth mammary fat pad of 6-8 weeks old female BALB/c mouse (SPF Biotechnology Co., Ltd, Beijing, China) [18]. To detect the compounds effects on the tumor growth and the mouse survival, 8 mg/kg/day of maraviroc (MedChemExpress, HY-13004) or verteporfin (TargetMol, T3112) was dissolved in PBS containing 2% DMSO and intraperitoneally injected to the mice daily [16]. Tumor growth was monitored daily with a caliper and tumor volume was calculated by the formula (mm^3^): length × width × height/2 [18]. Mice were sacrificed on the indicated day after injection and the bone marrow, spleens and tumors were harvested for further analysis. The humane endpoint for survival analysis of tumor bearing mice was setup by the professionals in the animal center of SJTU (Shanghai, China).

#### 2.2.2. TNBC tumor growth model in an immunodeficient mouse

##### 2.2.2.1. CD8^+^ T or Natural killer (NK) cells depleted mouse model

To deplete the CD8^+^ T cells in a mouse, 50 μg of anti-mouse CD8α (clone 2.43) antibody (BioXCell, BP0061) was intraperitoneally injected to the mouse on the days of -1^st^, 2^nd^, 7^th^, 12^th^, and 19^th^ post tumor inoculation [19] and the depletion efficiency of CD8^+^ T cells was verified by the fluorescence-activated cell sorting (FACS) analysis on day 18 post treatment.

To deplete the NK cells in a mouse, 20 μl of Rabbit anti-mouse/rat asialo GM1 polyclonal antibody (CEDARLANE, CL8955) was intravenously injected to the mouse on the days of -1^st^, 4^th^, and 9^th^ post tumor inoculation [20] and the depletion efficiency of NK cells was verified by the FACS analysis on day 12 post treatment.

2 × 10^5^ 4T1 cells were resuspended in 100 μl PBS and then injected into the fourth mammary fat pad of 6-8 weeks old CD8^+^ T cells depleted or NK cells depleted female BALB/c mouse. Tumor growth was monitored as described above and mice were sacrificed on the indicated day.

##### 2.2.2.2. *NOD-SCID* mouse *model*

2 × 10^5^ 4T1 cells were resuspended in 100 μl PBS and then injected into the fourth mammary fat pad of 6-8 weeks old female NOD-SCID mouse (Vital River Laboratory Animal Technology Co., Ltd, Beijing, China). Tumor growth was monitored as described above and mice were sacrificed on the indicated day.

#### 2.2.3. TNBC metastasis model

2 × 10^5^ 4T1 cells were resuspended in 200 μl of PBS and intravenously injected into the 6-8 weeks old female BALB/c mouse through the tail vein [21]. 3 weeks later, mice were sacrificed and the lungs were harvested for further analysis. To detect the macro-metastasis tumors, the harvest lungs were either immersed in the Bouin’s solution (including 75 ml of saturated picric acid, 25 ml of 4% paraformaldehyde and 5 ml glacial acetic acid) [22] and photographed, or stained with H&E. Since 4T1 is a thioguanine-resistant cell line, the 6-thioguanine assay was used to detect the micro-metastasis tumors [23].

### 2.3. Bioinformatics analysis

The data of mRNA expression of *CCL5* and *CCR5* in different cancers from The Cancer Genome Atlas (TCGA) and Genotype-Tissue Expression Project (GTEx) were analyzed by the Gene Expression Profiling Interactive Analysis website (GEPIA; http://gepia.cancer-pku.cn/index.html) [24]. The correlation of the expression of chemokines or chemokine receptors with cumulative survival of different cancer patients were evaluated by using the Tumor IMmune Estimation Resource website (TIMER; https://cistrome.shinyapps.io/timer/) [25]. The correlation of the expression of *CCL5* or *CCR5* with the infiltration level of Treg and MDSC cells in the tumor microenvironments in BRCA, GBM and ESCA was analyzed by using the TISIDB website (http://cis.hku.hk/TISIDB/) [26]. The correlation analysis of mRNA level between *CCR5* and *YAP1* in different cancers was performed by using the Tumor IMmune Estimation Resource 2.0 website (TIMER 2.0; http://timer.cistrome.org/) [27].

### 2.4. Construction of chemokines and chemokine receptors stable expressing cell lines

Human *CCL3* (P43969), *CCL4* (P18566), *CCR1* (P7780), *CCR3* (P6883) and *GPR75* (P35792) cDNAs were purchased from MiaoLing Plasmid Platform, human *CCL5*, *CCL7*, *CCL11* and *CCR5* cDNAs were gifts from Prof. Jiahuai Han (Xiamen University, Xiamen, China). The *CCL3*, *CCL4*, *CCL5*, *CCL7* and *CCL11* genes were designed with a C terminal 6 × His tag. All genes were subcloned into a pCDH-CMV-MCS-EF1-Puro vector plasmid and sequenced, respectively. The recombinant plasmids were packaged into the lentivirus with pSPAX2 (Addgene, 12260, RRID: Addgene_12260) and pMD2.G (Addgene, 12259, RRID: Addgene_12259) plasmids, and infected the HEK293T cells, an empty vector was infected as the control (HEK293T-EV). After selecting with 2 μg/ml of puromycin (Sigma-Aldrich, 540411) for a week, single clones were seeded, cultured and collected. The protein levels were detected by western blotting, respectively. The clone with the highest expression level was used for further experiments. Primers used for amplification of genes in this study are described in Supplementary Table S1.

### 2.5. Purification of recombinant human chemokines

Human chemokines stable expressing HEK293T cells were cultured in 15 cm dishes, the medium was collected every other day from the day 4 to day 28. The collected medium was then incubated with the Ni^+^ NTA beads 6FF (Smart-Lifesciences, SA005010) at 4°C overnight and transferred into the column. The beads were washed by 10 mM, 50 mM and 125 mM imidazole (Sigma-Aldrich, 288- 32-4), respectively, and then eluted by 250 mM imidazole. The purified chemokines were analyzed by SDS-polyacrylamide gel electrophoresis (SDS-PAGE) with silver staining, and also quantified by the Enhanced BCA Protein Assay Kit (Beyotime Biotechnology, P0009).

### 2.6. The high-throughput screening assay for discovering antagonists of CCL5/CCR5 axis

10,000 cells/well of human CCR5 stable expressing HEK293T cells were coated into a 96-well plate and cultured overnight. Cells were then fixed by 4% paraformaldehyde (Servicebio, G1101-500ML) for 20 min at room temperature (RT) in the dark and blocked with 5% BSA PBS buffer for 1 h at RT in a shaker. After washing with PBS buffer, compounds from different libraries were added into the well, and incubated for 3 h. 0.5% DMSO (Millipore, 196005) and 100 μM merbromin (Sigma-Aldrich, M7011) were used as a negative control or positive control, respectively. Then 30 μg/ml of purified His-tagged CCL5 was added into the wells and incubated for another 3 h at RT in the shaker, and the same volume of PBS buffer was added as a blank. The plate was then washed with the washing buffer (0.05% (v/v) Tween-20 PBS buffer) for three times and 100 μl of 1:5000 diluted HRP-conjugated anti-His antibody (Proteintech, HRP-66005) was added into each well and incubated at 4°C overnight or 2 h at RT. Plate was then washed with washing buffer for three times and 100 μl of ABTS solution (ThermoFisher Scientific, 002024) was added into the wells and reacted for 30 min in the dark. The products were then detected under O.D. 405 nm by a full-wavelength microplate reader (BioTek).

The Z-factor of the screening assay was calculated from 31 controls (0.5% DMSO) and 31 samples (100 μM merbromin) according to the below equation 1 [28].

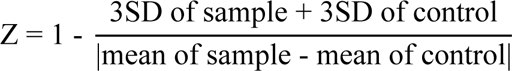

The inhibition effects of the compounds on CCL3, CCL4, CCL5, CCL7 or CCL11 binding to CCR5, as well as the CCL5 binding to GPR75, CCR1, CCR3 or CCR5, were also evaluated by this assay.

### 2.7. Confocal microscopy

Confocal microscopy was performed to detect the binding effects of different axes, and monitor the inhibition effects of the compounds on disrupting the interaction between the chemokine and the chemokine receptor.

For detecting the binding efficiency of chemokine to its receptor, cell slides were put into a 12-well plate and pre-treated with 0.1 mg/ml poly-D-lysine (ThermoFisher Scientific, A3890401). The chemokine receptor stable expressing HEK293T cells were then seeded into the slides and cultured overnight. After removing the supernatant and washing with PBS, cells were fixed by 4% paraformaldehyde for 20 min at RT and blocked with 5% BSA for 1 h in a shaker. 30 μg/ml purified chemokines were then added into the well. After washing with the washing buffer (0.05% (v/v) Tween-20 PBS buffer) for 10 min for three times, the anti-His antibody (Proteintech, 66005-1-Ig, 1:1,000) was added into the wells and incubated at 4℃ overnight. After washing as described above, the Alexa Fluor-488 conjugated goat anti-mouse IgG (H+L) antibody (Beyotime Biotechnology, A0428, 1:500) was added into the wells and incubated at RT in the dark for 2 h. After washing, DAPI (Beyotime Biotechnology, C1005) was added and incubated for 10 min in the dark without shaking. After washing, the slides were put in the antifade polyvinylpyrrolidone mounting medium (Beyotime Biotechnology, P0123-5 mL) and examined by an Upright Laser Confocal Microscope (Nikon Ni-E A1 HD25).

For monitoring the inhibition effects of the compounds on different axes, compounds were added into the wells and incubated for 3 h (0.5% DMSO were employed as the control) before adding the chemokines as described above in detecting the binding efficiency of chemokine to its receptor.

### 2.8. Cell migration assay

*In vitro* scratch assay was carried out to detect the cell migration [29]. Cells were seeded into a 6- well plate. Scratch was made on confluent monolayer in cell culture wells, control (0.1% DMSO) or compounds were added into each well. After 1 h, 30 μg/ml purified human CCL5 was added and incubated for 24 h for TNBC cells or 56 h for HCT-116 cells, respectively, the scratches were then documented by an Upright-Reverse Fluorescence Microscope (Revolve & ECHO), and the migrated distance was quantified using ImageJ software.

### 2.9. Cell invasion assay

To evaluate the invasive abilities of cancer cells, transwell chambers (Corning, 3422) with an 8-μm pore size were pre-coated with Matrigel (BD Biosciences, 354234) [30]. 1×10^5^ TNBC cells or 5×10^5^ HCT-116 cells were resuspended in 100 μl serum-free fresh medium and seeded into the upper chambers. And 600 μl growth medium containing 30 μg/ml purified CCL5 as well as different compounds was added to lower chamber, 0.1% DMSO was added as control. After TNBC cells invaded for 15 h or HCT-116 cells for 48 h, the cells that had invaded to the lower surface of the transwell membrane were fixed with 4% paraformaldehyde and stained with Crystal Violet Staining Solution (Beyotime Biotechnology, C0121-100 mL). Several random fields of views of this assay were captured by the Upright-Reverse Fluorescent Microscopy (Revolve & ECHO) and counted for each membrane.

### 2.10. 6-thioguanine assay

Lungs in the mice tumor metastasis model were excised, minced and digested by 1.5 mg/ml D- collagenase (Sigma-Aldrich, COLLD-RO) with periodic vortexing at 37°C for 1 h, and the erythrocytes were removed by red blood cell lysate (Sangon Biotech, B541001-0100). Cell suspensions were strained with 70 μm filters. The single-cell suspensions were then seeded in 6-well plates and cultured for 7 to 14 days in the presence of 60 μM 6-thioguanine (Sigma-Aldrich, A4882) [23], followed by fixing with 4% paraformaldehyde and staining by Crystal Violet Staining Solution. The number of colonies formed in each well was counted and photographed by ChemiDoc imaging system (DynaMax Biotech).

### 2.11. Construction of CCR5 and YAP1 knockdown cell lines

For generation of the *CCR5* knockdown cells, the TNBC cells were seeded in a 6-well plate at 70% to 80% confluence. A nontargeting siRNA scramble control (si*NC*) and siRNA targeting *CCR5* (si*CCR5*) were transfected into TNBC cells using lipofectamine^TM^ 2000 transfection reagent (ThermoFisher Scientific, 11668019) according to the manufacturer’s instructions. The cells were then harvested to detect the mRNA level 48 h post transfection and monitor the protein level 96 h post transfection.

For generation of the *YAP1* knockdown cell line, the *YAP1*-targeted shRNA 1 (sh*YAP1* #1), shRNA 2 (sh*YAP1* #2) or scramble shRNA (sh*NC*) were inserted into pLKO.1-TRC Cloning Vector (Addgene, 10878, RRID: Addgene_10878) following the manufacturer’s protocol and sequenced. The lentiviruses were generated by co-transfection of HEK293T cells with psPAX2, pMD2.G and the corresponding shRNA using Lipofectamine^TM^ 2000 reagent following the manufacturer’s instructions. The supernatants containing the lentivirus were collected and filtered through a 0.45 μm syringe filter 24 and 48 h post transfection. TNBC cells were then infected with the lentivirus for 24 h in the presence of polybrene (8 μg/ml) and selected with puromycin. The mRNA and protein level of YAP1 were detected by qPCR and western blotting, respectively.

The sequences of siRNAs or shRNAs used in this study are described in Supplementary Table S2.

### 2.12. Quantitative real-time PCR (qPCR)

Total RNA was extracted with TRIzol reagent (Invitrogen, 15596026) and cDNA was prepared with the HiScript II Q RT SuperMix for qPCR (+gDNA wiper) kit (Vazyme, R223-01). cDNA was mixed with Hieff^®^ qPCR SYBR Green Master Mix (Yeasen Biotechnology, 11201ES08) and subjected to qPCR with a sequence detection system (CFX; Bio-Rad Laboratories). The qPCR primers used in this study are described in Supplementary Table S3.

### 2.13. Western blotting

Cells were collected and lysed with RIPA (Beyotime Biotechnology, P0013B) supplemented with the protease and phosphatase inhibitor (Beyotime Biotechnology, P1048) and Phenylmethanesulfonyl fluoride (PMSF, Beyotime Biotechnology, ST506) for 30 min on ice and sonicated for 10 s under 50 amplitudes (ThermoFisher Scientific, FB705220). The lysates were then centrifuged at 12,000 × g for 10 min at 4°C, and supernatants were collected. Protein concentration was quantified by using the Enhanced BCA Protein Assay Kit. A total of 50 μg of proteins were separated by 10% or 15% SDS-polyacrylamide gel (SDS-PAGE) and transferred to a nitrocellulose membrane (Merck, HATF00010). To detect the CXCL16 and CXCL8 secreted in the culture medium, the same number of different cells were cultured in the 6-well plate for 48 h and the culture medium was collected, centrifuged and supernatants were separated by 10% or 15% SDS-PAGE and transferred to a nitrocellulose membrane. After blocking with 5% skim milk, the membranes were incubated first with the primary antibodies. Signal was then detected with fluorescent secondary antibodies. The membranes were finally detected by the LI-COR Odyssey CLx imaging system (LI-COR Biosciences) and quantified by normalization to GAPDH with ImageJ. The antibodies used in this study are described in Supplementary Table S4.

### 2.14. IHC analysis of the co-expression of CCR5 and YAP1 in human TNBC tumor samples

The co-expression of CCR5 and YAP1 in 10 female TNBC patient samples (aged from 32 to 60, including 4 in stage 1, 5 in stage 2, and 1 in stage 3, respectively. Among them, one patient was ipsilateral clavicle lymph node relapsed and none were metastasized) were analyzed by IHC. The sections from the same patient were detected with anti-CCR5 (Affinity Biosciences, AF6339, 1:300) or anti-YAP1 (Proteintech, 66900-1, 1:300) antibody, respectively. These experiments were performed and analyzed by Jilin Cancer Hospital (China).

### 2.15. MTS assay

Cells were seeded in 96-well plates at a density of 1 - 5 × 10^4^ cells per well. After the indicated treatments, cells in each well were incubated with 20 μl of MTS reagent (Promega, G3582) [31] at 37°C for 1-3 h. Cell viability was determined by measuring the absorbance of the resultant formazan dye at O.D. 490 nm by using a full-wavelength microplate reader (BioTek).

### 2.16. Cell apoptosis assay

Cell apoptosis was analyzed by using the Alexa Fluor^TM^ 488 Annexin V/Dead Cell Apoptosis Kit (ThermoFisher Scientific, V13241) [32] according to the manufacture’s protocol by flow cytometry. In brief, cells were collected, washed in cold phosphate-buffered saline (PBS), and suspended in 1 × annexin-binding buffer. Then, cells were stained with 5 μl of Alexa Fluor^TM^ 488 annexin V and 1 μl of propidium iodide at RT for 15 min in the dark, and then analyzed by flow cytometry.

### 2.17. TUNEL, Ki67, H&E and IHC staining

All specimens were fixed with 4% paraformaldehyde for 24 h and then embedded in paraffin, and sectioned at 3 μM. The sections were then deparaffinized at 60°C for 30 min, cleared in 100% xylene, and rehydrated with a graduated series of ethanol solutions from 100% to 70%.

For TUNEL and Ki67 staining, tumor sections were subjected to TUNEL and Ki67 staining using DAB (SA-HRP) Tunel Cell Apoptosis Detection Kit (Servicebio, G1507) or Ki67 antibody (Servicebio, GB111499, 1:500), respectively.

For H&E staining, the lung sections were stained with Hematoxylin-Eosin (HE) Stain Kit (Solarbio life sciences, G1120) according to the manufacturer’s protocols.

For IHC staining, tumor sections were treated with citrate buffer (pH 6.0) for antigen retrieval. After inhibition of endogenous peroxidase activity for 25 min with 3% H2O2, the sections were incubated with 3% BSA buffer for 30 min for blocking of non-specific binding. The sections were then incubated with the primary antibody at 4°C overnight and then incubated with the HRP-conjugated secondary antibody at RT for 1 h. The sections were further stained with DAB chromogenic reagent (DAKO, K5007) and counterstained with hematoxylin (Servicebio, G1004). All above experiments were performed by Servicebio Technology Co. Ltd. Wuhan. China. The antibodies used in this study are described in Supplementary Table S4.

At least five random fields of views per section were captured by the Upright Fluorescent Microscope (Axio Imager M2) and the percentage of apoptotic or proliferation cells, or CD4^+^, CD8^+^, CD49b^+^, Foxp3^+^, Ly6G^+^ cells were analyzed by ImageJ software.

### 2.18. Flow cytometry

The bone marrow cells were harvested from the femurs of tumor-bearing mice. The spleens were mashed and the erythrocytes were removed by red blood cell lysate. The tumors harvested from the tumor-bearing mice were digested with tissue dissociation buffer [D-collagenase (1 mg/ml), DNase I (50 μg/ml), and Dispase^®^ II (1 U/ml) in PBS] by periodic vortexing at 37°C for 1 h. Cell suspensions from the bone marrow, spleens and tumors were then strained with 70 μm filters. The single-cell suspensions were incubated in mouse Fc block (anti-CD16/32) antibody (Biolegend, 101335, 1:300) and then stained with the appropriate antibodies in FACS Buffer as described previously [33]. The antibodies used in this study are described in Supplementary Table S4.

### 2.19. ELISA

The mouse serum was collected in the indicated day from the tumor-bearing mice and were cold for 2 h on ice and then centrifuged at 1,000 × g at 4°C for 10 min, and supernatants were collected. The quantification of IL-10, TNF-α, IFN-γ and TGF-β in mouse serum was performed with the BD OptEIA^TM^ Mouse IL-10 ELISA Set (BD Biosciences, 555252), BD OptEIA^TM^ Mouse TNF (Mono/Mono) ELISA Set (BD Biosciences, 555268), BD OptEIA^TM^ Mouse IFN-γ ELISA Set (BD Biosciences, 555138) and Human/Mouse/Rat/Porcine/Canine TGF-beta 1 Quantikine ELISA (R & D Systems, DB100C) according to the manufacturer’s protocol, respectively.

### 2.20. RNA sequencing (RNA-seq) and data analysis

RNA was extracted from the control (si*NC*) and *CCR5* knockdown (si*CCR5*) MDA-MB-231 cells by using TRIzol reagent according to the manufacturers’ protocols. RNA-seq libraries were constructed by NEBNext^®^ Ultra RNA Library Prep Kit for Illumina (NEB, Cat# E7530L), and were sequenced by Illumina NovaSeq 6000 sequencing machine with 150-bp paired-end reads by Beijing Novel Bioinformatics Co., Ltd. (https://en.novogene.com/). The Raw reads were filtered using fastp (0.13.1) and mapped to the human reference genome (GRCh38/hg38) using HISAT2 (2.1.0) with parameters “--rna-strandness RF --dta”. Transcript abundance was normalized and measured by Fragments Per Kilobase of exon model per Million mapped fragments (FPKM). DESeq2 R package (1.20.0) was used to identify differential expression genes (DEGs). Genes with |log2FoldChange| > 1 and *p* < 0.05 were counted as DEGs. To identify functional categories of DEGs, KEGG and Gene Ontology enrichment analyses were performed by using the clusterProfiler R package (3.8.1). Pheatmap was used to display heatmaps of the expression levels of DEGs, which were normalized by z-score.

### 2.21. Statistical analysis

Data analysis was performed by using GraphPad Prism 6.0 or Excel 2019 software, and presented with error bars as mean ± SEM. One-way ANOVA and Unpaired two-tailed Student’s t test were used to compare experimental groups, and Kaplan–Meier method with the log-rank test was used for survival analysis. Statistical significance was defined as a *P* value of less than 0.05. Levels of significance were indicated as \**P* < 0.05; \*\**P* < 0.01; \*\*\**P* < 0.001; \*\*\*\**P* < 0.0001; ns, not significant. The statistical details including the statistical tests used, exact value of n, and precision measures were all specified in the figure legends unless otherwise indicated.

## 3. Results

### 3.1. Highly expressed CCL5 and CCR5 in intractable cancers are associated with poor prognosis

To identify the correlation of *CCL5* and *CCR5* with cancers, we analyzed the data from TCGA and GTEx by Gene Expression Profiling Interactive Analysis (GEPIA) and found that the mRNA expression of *CCL5* and *CCR5* are both significantly higher in tumor tissues than their normal counterparts in several intractable carcinomas such as BRCA (breast invasive carcinoma), GBM (glioblastoma multiforme) and PAAD (pancreatic adenocarcinoma) (Fig. 1A). High expression of *CCR5* is also significantly associated with poor cumulative survival of patients with BRCA, GBM, ESCA (esophageal carcinoma) and THYM (thymoma) (Fig. 1B). However, high expression of other ligands of CCR5 and other receptors of CCL5 are not significantly associated with poor cumulative survival of BRCA patients (Fig. S1A and B). In addition, expression of *CCL5* and *CCR5* presents strong and significant positive correlation with the infiltration level of immunosuppressive regulatory T cells (Tregs) (i) and myeloid-derived suppressor cells (MDSCs) (ii) in BRCA (Fig. 1C), GBM (Fig. 1D) and ESCA (Fig. 1E). These data suggest that the CCL5/CCR5 axis may play an important role in these cancers in a manner that involves immune cells in the tumor microenvironment (TME). Hence, discovery of antagonists of the CCL5/CCR5 axis could provide means for the development of efficient and immune therapeutic modalities.

**Fig. 1.**
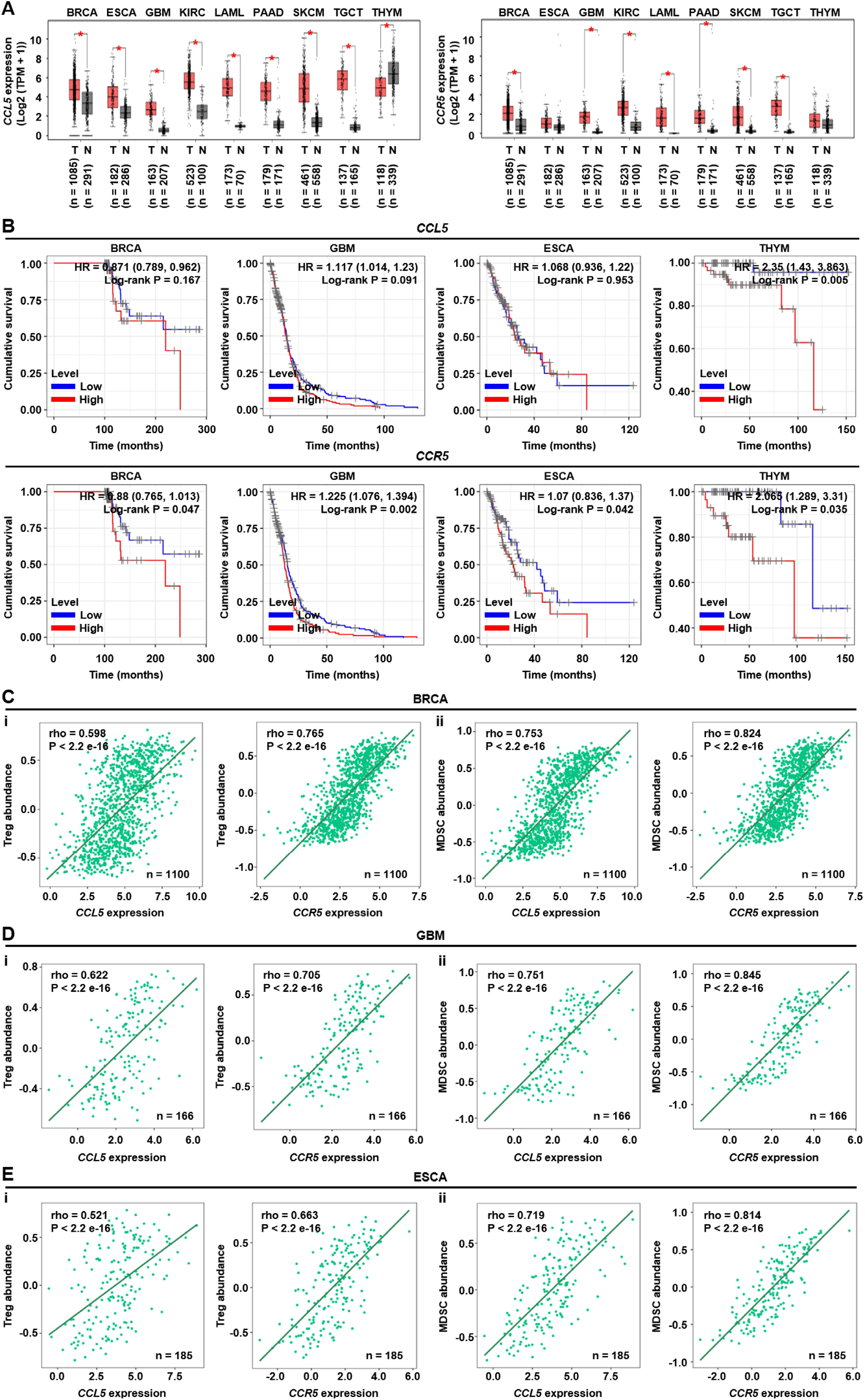
Highly expressed *CCL5* and *CCR5* in intractable cancers are associated with poor prognosis. (A) The expression of *CCL5* and *CCR5* in different tumor tissues from the TCGA database and paired normal tissues from the TCGA and GTEx databases were analyzed by the GEPIA website, respectively. TPM: Transcription per million, T: tumor, N: normal. BRCA: breast invasive carcinoma; ESCA: esophageal carcinoma; GBM: glioblastoma multiforme; KIRC: kidney renal clear cell carcinoma; LAML: acute myeloid leukemia; PAAD: pancreatic adenocarcinoma; SKCM: skin cutaneous melanoma; TGCT: testicular germ cell tumors; THYM: thymoma. (B) The cumulative survival of different cancers with high and low expression of *CCL5* and *CCR5* were analyzed by the TIMER website, respectively. (C-E) Correlations of *CCL5* and *CCR5* expression with the infiltration level of Tregs (i) and MDSCs (ii) in BRCA (C), GBM (D) and ESCA (E) from the TCGA were analyzed by the TISIDB website. Data in (A) was analyzed by one-way ANOVA. \**P* < 0.05.

### 3.2. A high-throughput screening assay for discovering antagonists of the CCL5/CCR5 axis

To identify specific antagonists of the CCL5/CCR5 axis, we established a cell-based high-throughput screening (HTS) assay in a 96 micro-well format in HEK293T cells stably expressing CCR5 (HEEK293T-CCR5) with added His-tagged recombinant CCL5 and the compounds to be screened (Fig. 2A). The CCR5-stable expressing cells (Fig. 2Bi) were coated in the wells of microplates. Then purified His-tagged CCL5 polypeptides (Fig. 2Bii and iii) were added to the wells, followed by the HRP-conjugated anti-His antibody to further amplify the binding signal, which yielded the dark green colored end products for colorimetric detection following the addition of the HRP substrate ABTS (Fig. 2C). However, with the strong inhibition of antagonists which were added before or simultaneously with chemokines, the binding of CCL5 to CCR5 could be disrupted, and the color product cannot be made even with the addition of ABTS. With the further optimization of the numbers of CCR5-stable expressing cells, the concentration of His-tagged CCL5 peptides and DMSO, the binding time of CCL5 with CCR5, assay buffer and detecting wave (Fig. S2A-F), the final assay condition was determined as follows: 10,000 CCR5 stable expressing cells, 30 μg/ml CCL5, 0.5% DMSO in PBS assay buffer, 3 h binding time and detecting under O.D. 405 nm. The Z-factor of the assay was also evaluated with equation 1, the value of 0.84 indicated that this assay was an excellent assay [28] for discovering the antagonists of CCL5/CCR5 axis (Fig. 2D).

**Fig. 2.**
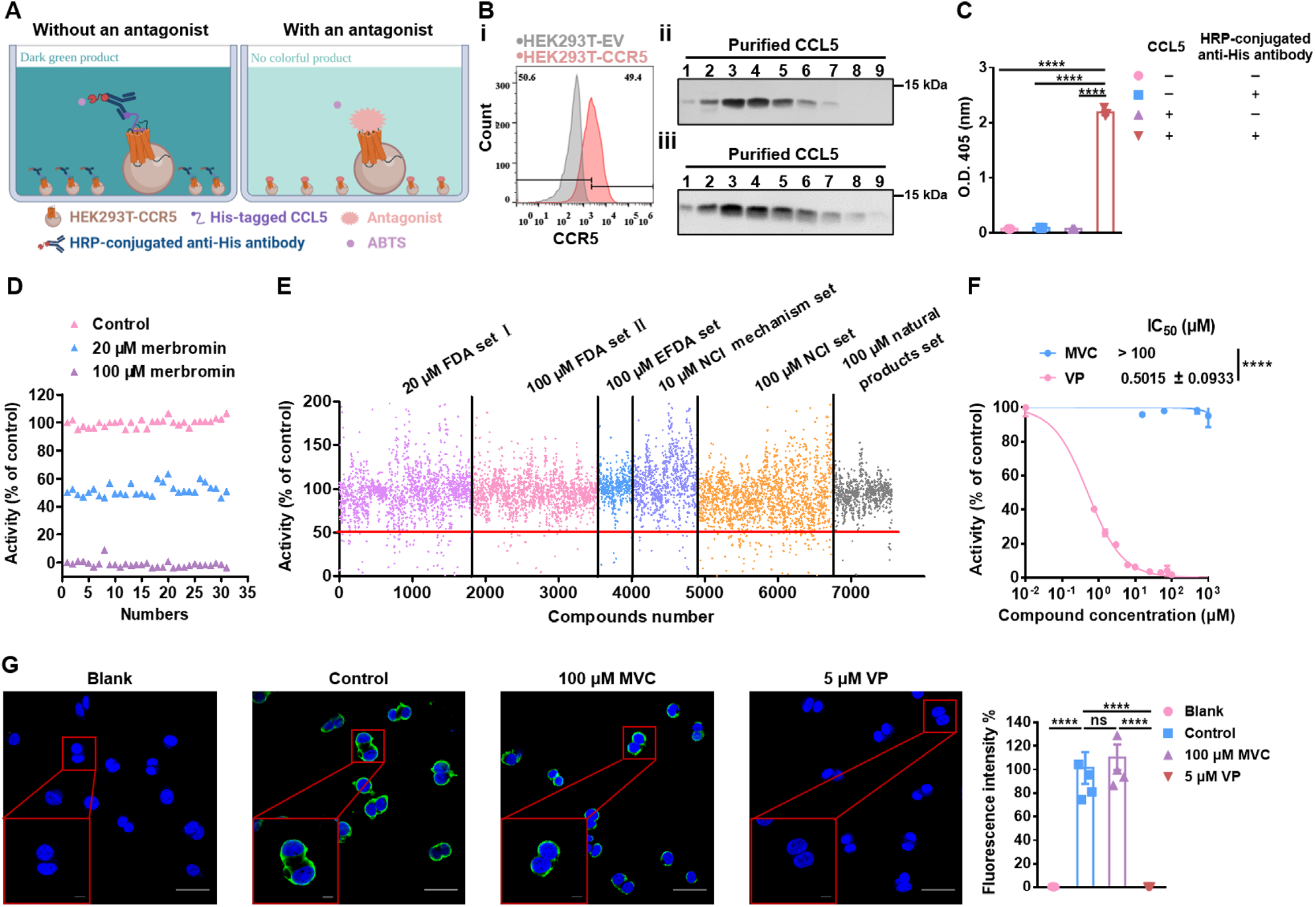
Establishing a high-throughput screening assay for discovering antagonists of the CCL5/CCR5 axis. (A) Schematic diagram of one micro-well of the developed cell-based high-throughput screening (HTS) assay for discovering the antagonists of CCL5/CCR5 axis. The CCR5- stable expressing cells (HEK293T-CCR5) were coated in the wells of a 96 micro-well plate overnight, after fixing with 4% paraformaldehyde for 20 min and blocking with 5% BSA for 1 h, the compound (antagonist candidate) was added into the well and incubated for 3 h, then the purified His-tagged CCL5 polypeptides were added and incubated for 3 h, after washing, the HRP-conjugated anti-His antibodies were then added and incubated for 2 h at room temperature (RT) or 4°C overnight, and the additional antibodies were then removed by washing. Finally, the substrate of HRP ABTS was added and detected under O.D. 405 nm in a microplate reader. Without an antagonist, the color of the well was in dark green, however, with an effective antagonist, the color was in light yellow green. (B) Characterization of the expression of CCR5 and the affinity purified His-tagged human CCL5 polypeptides. (i) The protein level of CCR5 on HEK293T-EV (the empty vector was transfected into HEK293T cells) and HEK293T-CCR5 cells were detected by flow cytometry. (ii-iii) The purified His-tagged CCL5 was analyzed by SDS-PAGE with silver staining (ii), and detected with the anti-His antibody by western blotting (iii). 1-9: The purified CCL5 by affinity purification. (C) The signal specificity analysis of the HTS assay was detected under O.D. 405 nm with or without addition of CCL5 or HRP-conjugated anti-His antibody (n = 3). (D) Analysis of the well-to-well reproducibility of the developed HTS assay in the 96 micro-wells. 0 (Control; ▲), 20 (▲) and 100 μM (▲) merbromin was added as the compounds in the assay described in above (A), respectively (n = 31). (E) High-throughput screening for identifying the antagonists of CCL5/CCR5 axis from 7,555 compounds, including 1,840 compounds (20 μM) from FDA set I, 1,700 compounds (100 μM) from FDA set II, 480 compounds (100 μM) from EFDA set, 879 compounds (10 μM) from NCI mechanism set, 1,856 compounds (100 μM) from NCI set and 800 compounds (100 μM) from natural products set. (F) IC50 analysis of maraviroc (MVC) and verteporfin (VP) on inhibition of the binding of CCL5 to CCR5 (n = 3). (G) The inhibition effects of MVC and VP on disrupting the binding of CCL5 to CCR5 was analyzed by confocal microscopy and quantified (n = 5) by ImageJ. Blank: without CCL5; Control: with 30 μg/ml of CCL5. Data in (C) and (G) were analyzed by one-way ANOVA, and data in (F) was analyzed by Unpaired two-tailed Student’s t test. Data were reported as mean ± SEM. \*\*\*\**P* < 0.0001, ns: not significant.

Using this system, we evaluated a total of 7,555 compounds from the FDA set I (1,840 compounds), FDA set II (1,700 compounds), EFDA set (480 compounds), NCI mechanism set (879 compounds), NCI set (1,856 compounds) and natural products set (800 compounds), respectively, for their ability to block CCL5 binding to CCR5 (Fig. 2E). Out of the numerous potential blockers, verteporfin was identified as a specific antagonist of the CCL5/CCR5 interaction with an IC50 of 0.5015 ± 0.0933 μM, which was at least 200 times greater than the FDA approved antagonist of CCR5, maraviroc (Fig. 2F). The differential capacity of verteporfin versus maraviroc was also observed in that the green fluorescence derived from the binding of CCL5 to CCR5, as analyzed by confocal microscopy. The green fluorescence was only weakly decreased by 100 μM maraviroc, but almost completely abolished with 5 μM verteporfin (Fig. 2G). Hence, based on the HTS assay, verteporfin is a far more superior antagonist than maraviroc for disrupting the CCL5/CCR5 axis.

### 3.3. Verteporfin is a specific probe for CCL5/CCR5 axis

Verteporfin has been reported to inhibit YAP1/TAZ-TEA domain family member 1 (TEAD) transcriptional activity, cell proliferation and CD44 expression in gastric cancer stem cells [34]. Thus, we compared the potency of verteporfin in blocking the CCL5/CCR5 interaction with two other known inhibitors of YAP1, CA3 and TED-347 [35, 36]. Fig. 3A showed that the IC50 of both CA3 and TED- 347 were over 100 μM, much higher than that of verteporfin at about 0.5 μM, indicating that verteporfin was a potent antagonist of the CCL5/CCR5 interaction, whereas the other YAP1 inhibitors were not.

**Fig. 3.**
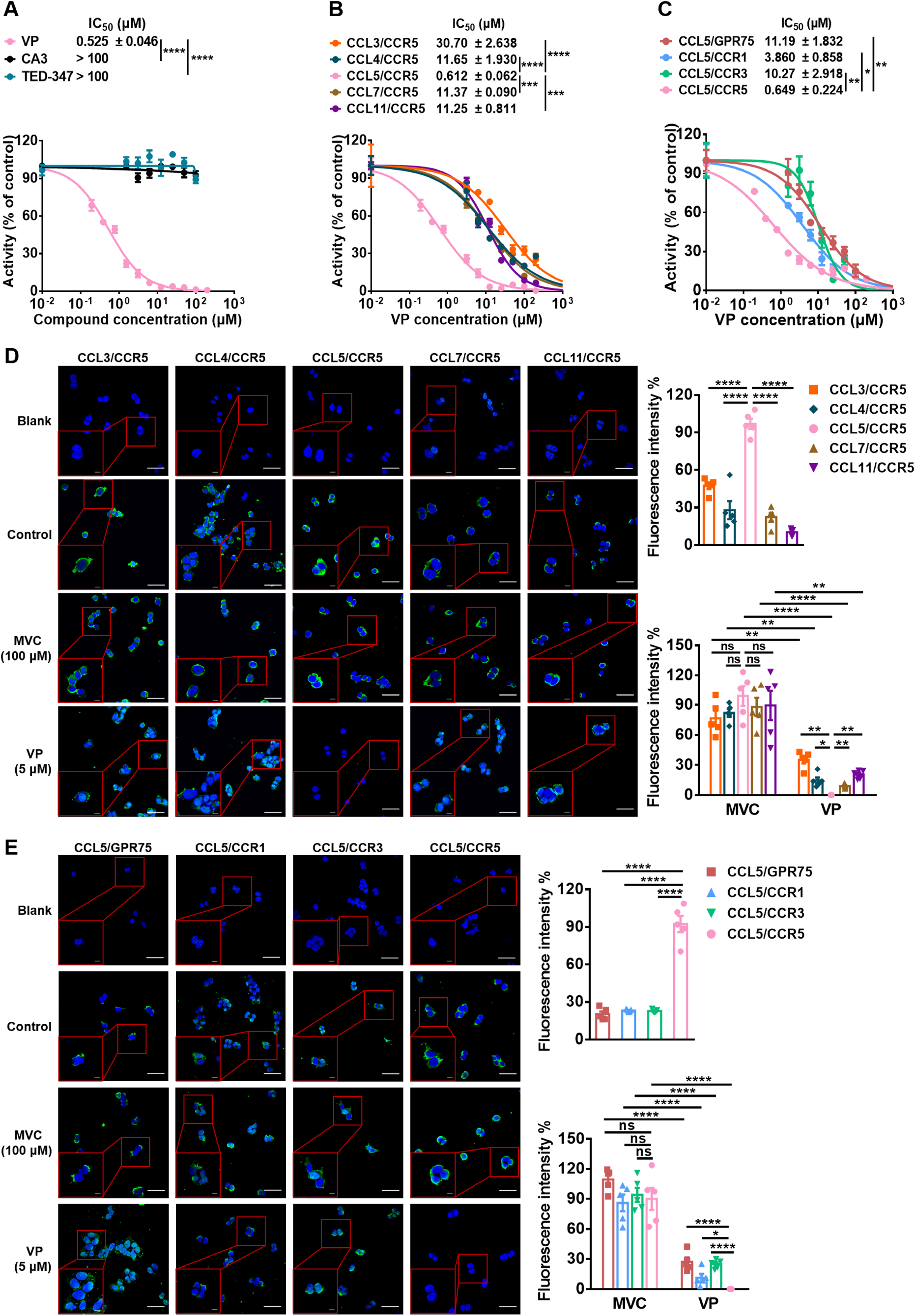
Verteporfin is identified as a specific probe for CCL5/CCR5 axis. (A) IC50 analysis of three YAP1 inhibitors verteporfin (VP), CA3 and TED-347 on inhibition of the binding of CCL5 to CCR5 (n = 3). (B-C) IC50 analysis of VP on inhibiting the binding of different chemokines of CCL3, CCL4, CCL5, CCL7 and CCL11 to CCR5 (B) or CCL5 binding to different receptors of GPR75, CCR1, CCR3 and CCR5 (C) (n = 3). (D) The binding effects of different chemokines (CCL3, CCL4, CCL5, CCL7 and CCL11) to CCR5, and the inhibition effects of maraviroc (MVC) and VP on these bindings were analyzed by confocal microscopy and quantified (n = 5) by ImageJ. Blank: without chemokines, Control: with 30 μg/ml of chemokines. (E) The binding effects of CCL5 to different receptors (GPR75, CCR1, CCR3 and CCR5) and the inhibition effects of MVC and VP on these bindings were analyzed by confocal microscopy and quantified (n = 5) by ImageJ. Blank: without CCL5, Control: with 30 μg/ml of CCL5. Data in (A-E) were analyzed by one-way ANOVA. Data are presented as mean ± SEM. \**P*< 0.05, \*\**P* < 0.01, \*\*\**P* < 0.001, \*\*\*\**P* < 0.0001, ns: not significant.

In addition to CCL5, C-C chemokine ligand 3 (CCL3), C-C chemokine ligand 4 (CCL4), C-C chemokine ligand 7 (CCL7) and C-C chemokine ligand 11 (CCL11) also interact with CCR5 [37, 38]. Conversely, G protein-coupled receptor 75 (GPR75), C-C chemokine receptor 1 (CCR1) and C-C chemokine receptor 3 (CCR3) are other receptors for CCL5 [39-41]. Thus, we further investigated the disrupting effects of verteporfin on other axes by both IC50 measurement and confocal microscopy analysis. The IC50 of verteporfin was about 0.6 μM on the CCL5/CCR5 axis, which was significantly lower than the CCL3/CCR5, CCL4/CCR5, CCL7/CCR5, and CCL11/CCR5 axis (Fig. 3B), as well as the CCL5/GPR75, CCL5/CCR1, and CCL5/CCR3 axis (Fig. 3C). Moreover, we compared the inhibition effects of maraviroc and verteporfin on these axes by confocal microscopy and confirmed that verteporfin was a much stronger disruptor of these axes than maraviroc (Fig. 3D and E). Taken together, compared to maraviroc, verteporfin is a specific probe for the CCL5/CCR5 axis.

### 3.4. Verteporfin significantly inhibits TNBC tumor growth without photodynamic treatment (PDT) mainly via immune effects

Verteporfin is known as a photodynamic reagent for reducing TNBC tumor growth and metastasis [42, 43]. Without light activation, verteporfin was found to inhibit cell proliferation in subtypes of breast cancer cell lines [44]. However, without PDT, the effects and mechanisms of verteporfin-mediated inhibition of TNBC tumor growth *in vivo* remain obscure. Here we evaluated them *in vitro* and *in vivo*. *In vitro*, compared to maraviroc, verteporfin treatment significantly decreased cell viability in the human ER^+^ breast cancer cell line MCF-7, the human TNBC cell lines of MDA-MB-231 and BT549, and murine TNBC cells of 4T1, as well as colon cancer cells of HCT-116 (Fig. 4A). Notably, compared to cisplatin (DDP), verteporfin treatment also reduced the viability of DDP-resistant MDA-MB-231 cells at the concentration of 6.25 μM or above, whereas DDP treatment at this dose did not (Fig. 4B). In addition, compared to maraviroc, verteporfin treatment significantly induced the apoptosis of MDA-MB-231, BT549 and 4T1 cells at the concentration of 5 μM or above for MDA-MB-231, and 0.6 μM for BT549 and 4T1 cells, respectively (Fig. 4C-E and Fig. S3A-C).

**Fig. 4.**
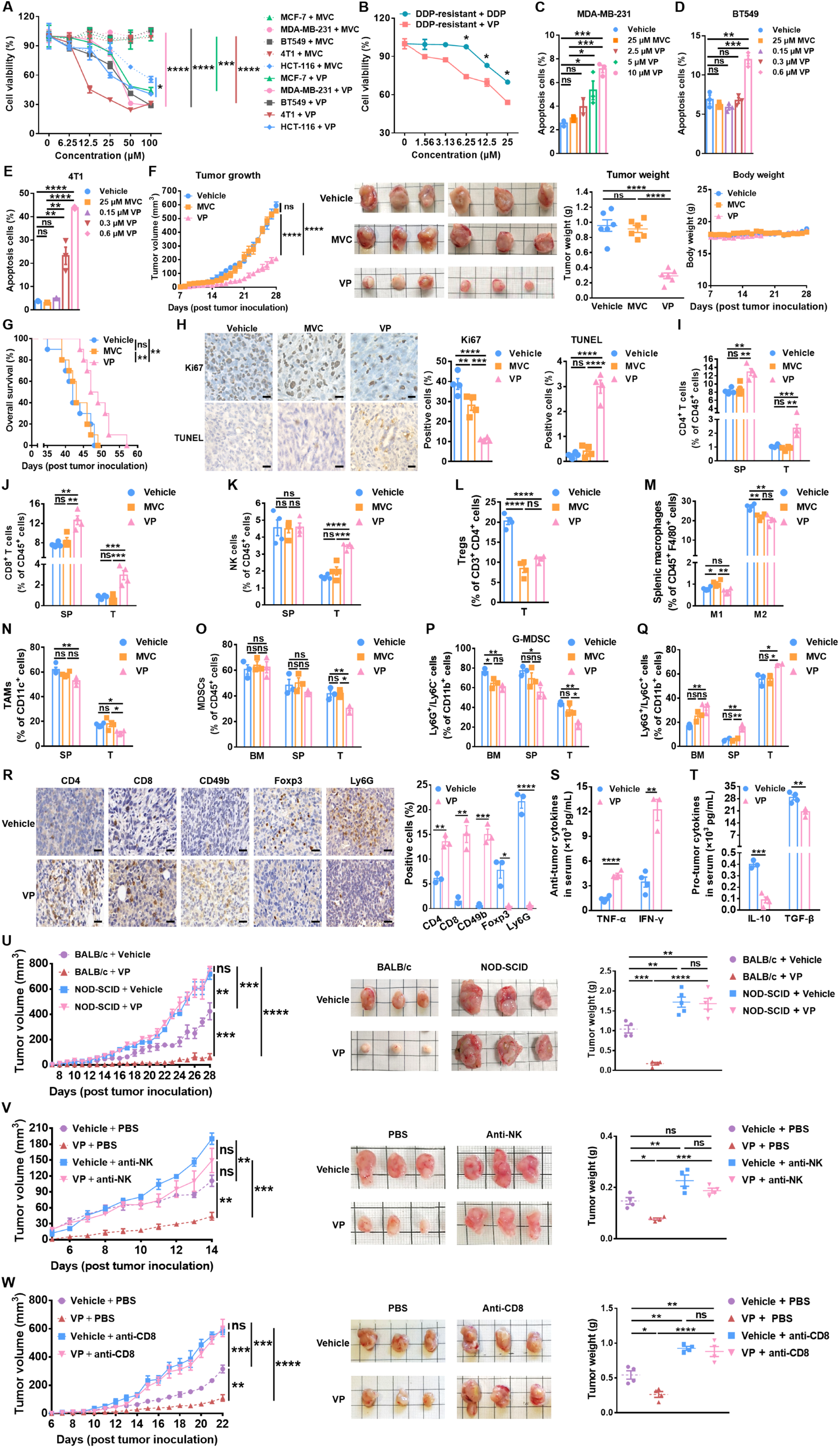
Verteporfin inhibits TNBC tumor growth without PDT mainly via immune effects. (A) The cell viability was measured in the human ER^+^ breast cancer cell line MCF-7, the human TNBC cell lines of MDA-MB-231 and BT549, and murine TNBC cells of 4T1, as well as human colon cancer cells of HCT-116 by MTS assay with maraviroc (MVC) or verteporfin (VP) treatment, respectively (n = 3). (B) The cell viability was measured in cisplatin (DDP)-resistant MDA-MB-231 cells by MTS assay with 0, 1.56, 3.13, 6.25, 12.5 and 25 μM of DDP or VP treatment, respectively (n = 3). (C-E) The quantification of apoptosis analyzed by flow cytometry in TNBC cells of MDA-MB-231 (C), BT549 (D) and 4T1 (E) treated with 25 μM MVC or indicated concentrations of VP, respectively (n = 3). (F) The tumor size and the body weight of mice were measured daily in BALB/c mice injected with 2×10^5^ 4T1 cells into the mammary fat pad and treated with vehicle, 8 mg/kg/day of MVC or VP, respectively. The representative tumor images were documented and the tumor weight was measured when the tumor-bearing mice were sacrificed on day 28 post injection (n = 6). (G) Kaplan-Meier analysis of the overall survival of the 4T1 tumor-bearing mice treated with vehicle, MVC or VP, respectively, as described in (F). One point denotes one mouse (n = 10). (H) Representative images and quantification of Ki67 and TUNEL staining of the tumor sections analyzed by IHC on day 28 post injection (n = 4). Scale bar: 10 μm. (I-K) The CD4^+^ T cells (I), CD8^+^ T cells (J) and NK cells (K) in spleens (SP) and tumors (T) of 4T1 tumor-bearing mice treated with vehicle, MVC or VP as described in (F) were analyzed by flow cytometry on day 14 post injection (n = 4). (L-N) The Tregs in T (L), macrophages in SP (M) and TAMs in SP and T (N) of 4T1 tumor-bearing mice treated with vehicle, MVC or VP as described in (F) were analyzed by flow cytometry on day 14 post injection (n = 4). (O- Q) Total MDSCs (O), G-MDSCs (P), and Ly6G^+^/Ly6C^+^ MDSCs (Q) in bone marrow (BM), SP and T of 4T1 tumor-bearing mice treated with vehicle, MVC or VP as described in (F) were analyzed by flow cytometry on day 28 post injection (n = 3). (R) Representative images and quantification of CD4^+^, CD8^+^, CD49b^+^ and Foxp3^+^ cells for the tumor sections analyzed by IHC on day 14, as well as Ly6G^+^ cells on day 28 post injection (n = 3). Scale bar: 10 μm. (S-T) Serum anti-tumor cytokines (S) and pro-tumor cytokines (T) of 4T1 tumor-bearing mice treated with vehicle or VP as described in (F) were analyzed by ELISA on day 14 post injection (n = 4). (U) The tumor size was measured daily in immune competent BALB/c (n = 4) or immunodeficient NOD-SCID (n = 5) mice treated with vehicle or 8 mg/kg/day of verteporfin (VP), respectively. The representative tumor images were documented and the tumor weight was measured when the tumor-bearing mice were sacrificed on day 28 post tumor inoculation. (V) The tumor size was measured daily in 4T1 tumor-bearing BALB/c mice treated with or without anti-mouse/rat asialo GM1 (anti-NK) antibody, or combined with vehicle or 8 mg/kg/day of VP, respectively. The representative tumor images were documented and the tumor weight was measured when the tumor-bearing mice were sacrificed on day 14 post tumor inoculation (n = 4). (W) The tumor size was measured daily in 4T1 tumor-bearing BALB/c mice treated with or without anti-CD8α (anti-CD8) antibody, or combined with vehicle or 8 mg/kg/day of VP, respectively. The representative tumor images were documented and the tumor weight was measured when the tumor-bearing mice were sacrificed on day 22 post tumor inoculation (n = 4). The square in the tumor image is 1cm × 1cm. Data in (A, B, and R-T) were analyzed by Unpaired two-tailed Student’s t test, data in (C-F, H-Q, and U-W) were analyzed by one-way ANOVA, and data in (G) was analyzed by Kaplan– Meier method with the log-rank test. Data are presented as mean ± SEM. \**P* < 0.05, \*\**P* < 0.01, \*\*\**P* < 0.001, \*\*\*\**P* < 0.0001, ns: not significant.

In the immune competent mouse 4T1 TNBC model, compared to maraviroc, verteporfin treatment without PDT markedly reduced the tumor growth and tumor weight, while the body weight of the mice was not affected (Fig. 4F). Verteporfin treatment also yielded a significant increase in overall survival (Fig. 4G). In addition, compared to vehicle, both maraviroc and verteporfin treatment decreased the number of Ki67^+^ cells in tumor sections. However, only verteporfin treatment increased TUNEL-positive apoptotic tumor cells (Fig. 4H). Notably, an increase in infiltrating CD4^+^ and CD8^+^ T cells were detected by flow cytometry only in the verteporfin-treated group but not the maraviroc-treated group in both spleens and tumors (Fig. 4I and J, and Fig. S3D and E), while an increase in tumor-infiltrating NK cells was also detected in verteporfin-treated mice (Fig. 4K and Fig. S3F). Simultaneously, except the intratumoral Tregs and the splenic M2 macrophage (Fig. 4L and M), the splenic and intratumoral tumor-associated macrophages (TAMs; Fig. 4N), and MDSCs (Fig. 4O and Fig. S3G) were significantly decreased by the verteporfin treatment but not by the maraviroc treatment. Decreased Ly6G^+^/Ly6C^-^ MDSCs (G-MDSC) and increased Ly6G^+^/Ly6C^+^ MDSCs were observed in the bone marrow, spleens and tumors with the verteporfin treatment (Fig. 4P and Q, and Fig. S3H). IHC analyses of the intratumoral CD4^+^, CD8^+^, CD49b^+^, Foxp3^+^ and Ly6G^+^ cells (Fig. 4R), as well as the anti-tumor cytokines of TNF-α and IFN-γ, and the pro-tumor cytokines of IL-10 and TGF-β in mice serum (Fig. 4S and T) were also consistent with the above immune cells monitored, suggesting that verteporfin treatment had a strong impact on the TME in favor of anti-tumor immune responses.

To establish the immunological role of verteporfin treatment, we performed an *in vivo* tumor growth study in immunodeficient NOD-SCID mice. Compared to the tumor growth in immune competent BALB/c mice in which 4T1 tumor growth was strongly inhibited by the verteporfin treatment, there was no difference in the tumor growth and tumor weight in NOD-SCID mice with or without verteporfin treatment (Fig. 4U). This result confirmed that the effects of verteporfin on tumor growth were mediated mainly through its impact on immunological activities in TME.

To further assess the effects of verteporfin on the various immune populations in TME without PDT, we used specific antibodies to deplete NK and CD8^+^ T cells in the 4T1 model treated with verteporfin. The depletion efficiency of anti-NK and anti-CD8 antibodies were confirmed by flow cytometry (Fig. S4A and B). Both anti-NK and anti-CD8 antibody treatments strongly reversed the benefit of verteporfin treatment in controlling tumor growth and tumor weight (Fig. 4V and W). It is interesting to note that, the significant increases of Ly6G^+^/Ly6C^+^ MDSCs in bone marrow, spleens and tumors, as well as the significant decrease of G-MDSCs and MDSCs in tumors in the verteporfin alone treatment group, were not observed in the combination treatment of verteporfin with anti-NK antibody (Fig. S4C and D). In addition, compared to the verteporfin single treatment, the immunosuppressive cells of MDSC and G-MDSC cells in tumor were also increased, and the Ly6G^+^/Ly6C^+^ MDSC cells in tumor were decreased in the combination treatment of verteporfin with anti-CD8 antibody (Fig. S4E and F), resulted in the increase of tumor growth (Fig. 4V and W). These results suggested that verteporfin decreased TNBC tumor growth mainly by immune effects in TME.

### 3.5. CCR5 is essential for YAP1 expression and mediates a leukocyte activation program via YAP1

In MDA-MB-231 cells, knocking down *CCR5* expression via siRNA markedly decreased both mRNA and protein expression of YAP1, whereas knocking down *YAP1* expression via short hairpin RNAs (shRNAs) did not affect the expression of CCR5 in either mRNA or protein (Fig. 5A and B), indicating that CCR5 was essential for YAP1 expression. To explore the potential cell-intrinsic and extrinsic impacts of CCR5-mediated signaling in tumor cells, we performed an RNA-seq analysis using MDA-MB-231 cells with or without siRNA targeting of *CCR5*, and the resulting data were analyzed by Kyoto Encyclopedia of Genes and Genomes (KEGG) pathway analysis of the differentially expressed genes (DEGs). Compared to the control (si*NC*), a total of 381 genes were up-regulated and 939 genes were down-regulated in *CCR5* knockdown cells (si*CCR5*; Fig. 5C). The up-regulated genes were enriched in pathways such as cell cycle, DNA replication and hippo signaling pathway, whereas the down-regulated were enriched in IL-17, TNF, PI3K-Akt, and NF-kappa B signaling pathways (Fig. 5D). Interestingly, an analysis of the data by Gene Ontology (GO) terms for DEGs revealed an enrichment in leukocyte migration program, including the up-regulated chemokine C-X-C chemokine ligand 16 (*CXCL16*) and the down-regulated chemokine C-X-C chemokine ligand 8 (*CXCL8*, *IL-8*) in the si*CCR5* group (Fig. 5E and F), and they were among the top up-regulated or down-regulated DEGs potentially involved in leukocyte migration. Thus, we then further confirmed their differential gene expression regulated by CCR5 and YAP1 by knocking down technique and qPCR (Fig. 5G), as well as the protein level by western blotting (Fig. 5H). Since colon and breast cancer cell-expressed CXCL16 has been reported to bind to CXCR6 on Th1, NK, CD4^+^ T and activated CD8^+^ effector T cells and recruit them to the sites of inflammation [45-47], and tumor-produced CXCL8 helps the recruitment of immunosuppressive MDSCs, TAMs and Tregs [48-50], these results were consistent with our findings that tumor-infiltrating CD4^+^ T, CD8^+^ T and NK cells were increased, whereas immunosuppressive cells such as MDSCs, TAMs and Tregs were decreased in TME (Fig. 4I-K, L, N, and O), suggesting that tumor-derived CCR5/YAP1 may influence the migration of immune and immunosuppressive cells extrinsically via the chemokines of CXCL16 and CXCL8.

**Fig. 5.**
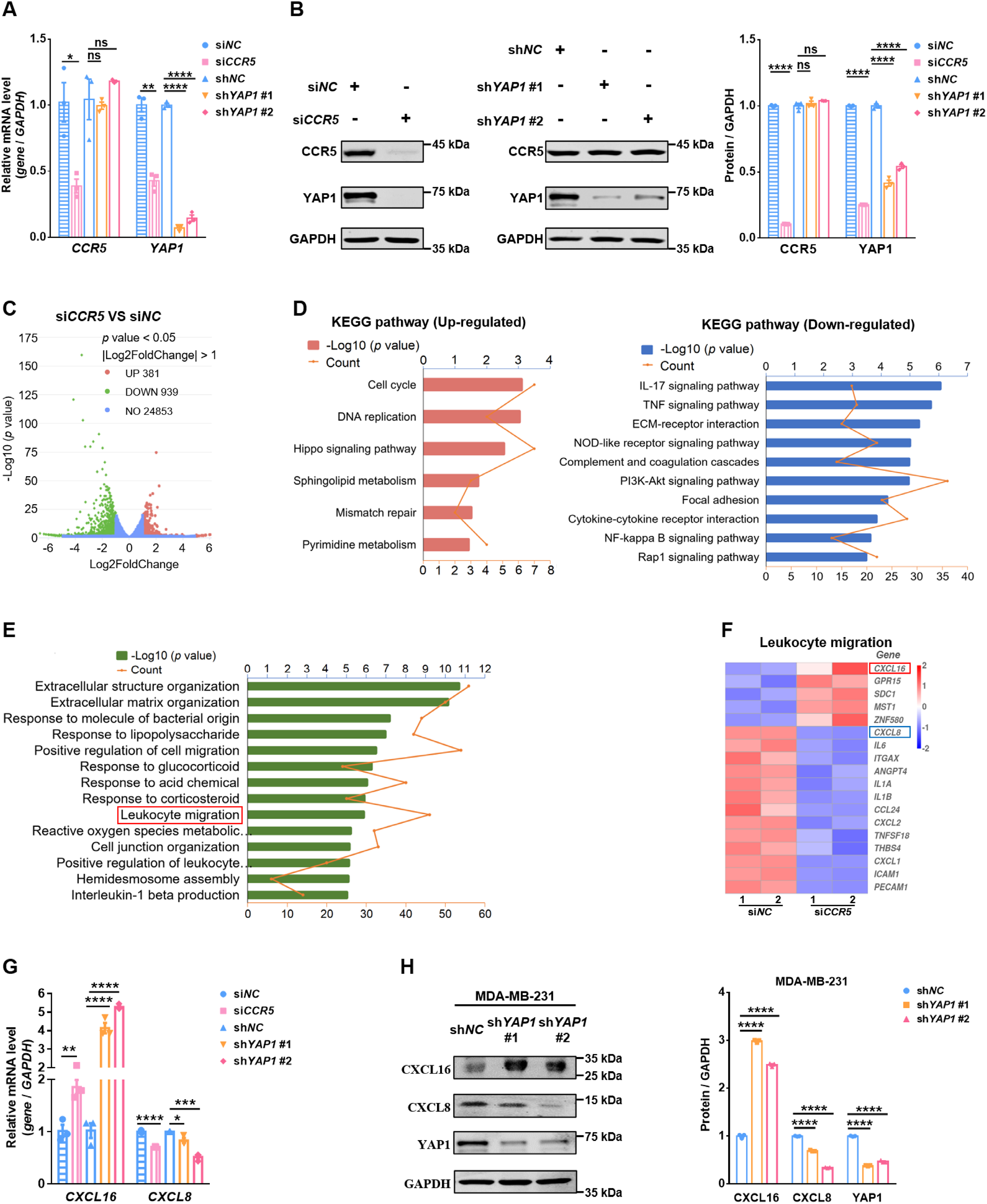
CCR5 is essential for YAP1 expression and mediates a leukocyte activation program via YAP1. (A-B) The mRNA expression of *CCR5* and *YAP1* was measured by qPCR (A) and their protein level was analyzed by western blotting (B) in control (si*NC*) and *CCR5* knockdown (si*CCR5*), as well as control (sh*NC*) and *YAP1* knockdown (sh*YAP1* #1 and sh*YAP1* #2) MDA-MB-231 cells (n = 3). (C) Volcano plot showing differentially expressed genes (DEGs) in si*NC* and si*CCR5* MDA-MB-231 cells identified by RNA-seq. (D) KEGG pathway analysis of DEGs in si*NC* and si*CCR5* MDA-MB-231 cells. Data were analyzed by using the clusterProfiler R package (3.8.1). (E) Enrichment of DEGs in si*NC* and si*CCR5* MDA-MB-231 cells for Gene Ontology terms. (F) Heatmap of the top 5 up-regulated and the top 13 down-regulated genes involved in leukocyte migration in si*NC* and si*CCR5* MDA-MB- 231 cells. (G) The mRNA level of *CXCL16* and *CXCL8* were measured by qPCR in si*NC*, si*CCR5*, sh*NC*, sh*YAP1* #1 and sh*YAP1* #2 MDA-MB-231 cells (n = 3). (H) The protein level of CXCL16, CXCL8 and YAP1 were monitored by western blotting in sh*NC,* sh*YAP1* #1 and sh*YAP1* #2 MDA-MB-231 cells (n = 3). Data in (A, B, G, and H) were analyzed by One-way ANOVA. Data are presented as mean ± SEM. \**P* < 0.05, \*\**P* < 0.01, \*\*\**P* < 0.001, \*\*\*\**P* < 0.0001, ns: not significant.

### 3.6. Verteporfin markedly reduces tumor cell metastasis without PDT

For investigation of the effects of verteporfin on tumor cell movement without PDT, we carried out migration assay and invasion assay with select cancer cell lines. Compared to the control, both maraviroc and verteporfin treatment significantly decreased the migratory and invasive capacities of TNBC cell lines of MDA-MB-231, BT549 and 4T1, and colon cancer cell line HCT-116 (Fig. 6A and B). Verteporfin was much more efficient than maraviroc in inhibiting cell migration (Fig. 6A) and invasion (Fig. 6B) with 5 to 10 times lower concentrations measured by the scratch assay and transwell invasion assay, in the absence of PDT. Moreover, in the 4T1 TNBC metastasis mouse model of tail vein injection, verteporfin treatment was considerably more potent than maraviroc in inhibiting lung metastasis as measured by the number of metastatic nodules in the lungs (Fig. 6C), tumor cell colonies grown in culture dishes in the presence of 6-thioguanine (Fig. 6D) and metastatic foci in the lungs visualized by Hematoxylin-Eosin (HE) Stain Kit (Fig. 6E). Hence, compared to maraviroc, verteporfin is a much more potent antagonist for tumor cell migration and metastasis without PDT.

**Fig. 6.**
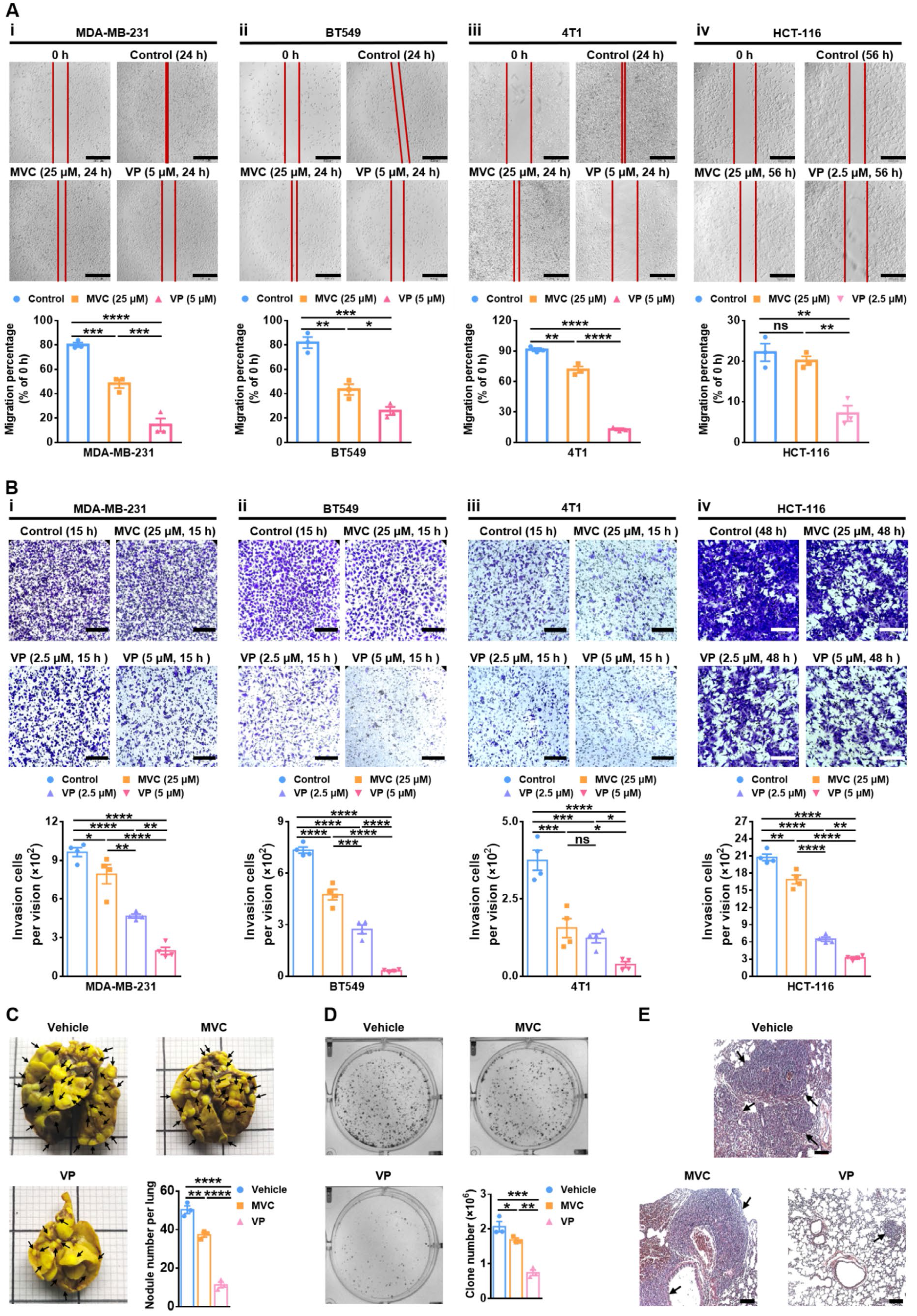
Verteporfin reduces tumor metastasis without PDT. (A) Representative images and quantification of scratch assay of control (0.1% DMSO), 25 μM maraviroc (MVC) and 5 μM verteporfin (VP) treated for TNBC cells of MDA-MB-231 (i), BT549 (ii) and 4T1 (iii), and 2.5 μM VP treated for colon cancer cells of HCT-116 (iv) (n = 3). Scale bar: 520 μm. (B) Representative images and quantification of transwell invasion assay of control (0.1% DMSO), 25 μM MVC, 2.5 μM and 5 μM VP treated for TNBC cells of MDA-MB-231 (i), BT549 (ii) and 4T1 (iii), and colon cancer cells of HCT-116 (iv) (n = 4). Scale bar: 220 μm. (C) Representative images and quantification (n = 3) of the visible metastatic nodules on lung surfaces in the 4T1 lung metastasis mice. Female BALB/c mice were intravenously injected with 2 × 10^5^ 4T1 cells and treated with vehicle, 8 mg/kg/day of MVC or VP, respectively. The tumor-bearing mice were sacrificed and the lungs were harvested on day 21 post injection and infused with Bouin’s solution. The arrows denote the metastatic nodules. The square in the lung image is 1cm × 1cm. (D) Metastatic tumor cells of the lungs as described in (C) were visualized and quantitated (n = 3) by the 6-thioguanine clonogenicity assay. (E) Representative H&E- stained images of lung metastatic foci. Scale bar: 40 μm. Data in (A-D) were analyzed by One-way ANOVA. Data are presented as mean ± SEM. \**P* < 0.05, \*\**P* < 0.01, \*\*\**P* < 0.001, \*\*\*\**P* < 0.0001, ns: not significant.

### 3.7. CCR5 is a key target of verteporfin and promotes metastasis via YAP1

Using the *CCR5* and *YAP1* knocking down MAD-MB-231 cells (Fig. 5A and B), decreased cell viability and increased apoptosis were observed by knocking down either *CCR5* or *YAP1* in MDA-MB-231 cells (Fig. 7A-C), although there were subtle differences in the amount of early versus late apoptosis caused by *CCR5*- or *YAP1*-expression silencing, respectively (Fig. 7C). In addition, the migratory and invasive abilities of MDA-MB-231 cells were also significantly decreased in *CCR5* knockdown cells (si*CCR5* MDA-MB-231) compared to their respective controls (si*NC* MDA-MB- 231), however, there was no significant difference for them when compared to the verteporfin treated control cells (Fig. 7Di and Ei, and Fig. S5Ai and Bi). Interestingly, the addition of verteporfin in the *YAP1* knockdown cells (sh*YAP1* MDA-MB-231) could further increase the inhibition of the migratory and invasive abilities of cells, whereas not for *CCR5* knockdown cells, suggesting that the metastasis driven by CCR5 only partially via YAP1, there were still other pathways involved in CCR5 mediated metastasis (Fig. 7Dii and Eii, and Fig. S5Aii and Bii).

**Fig. 7.**
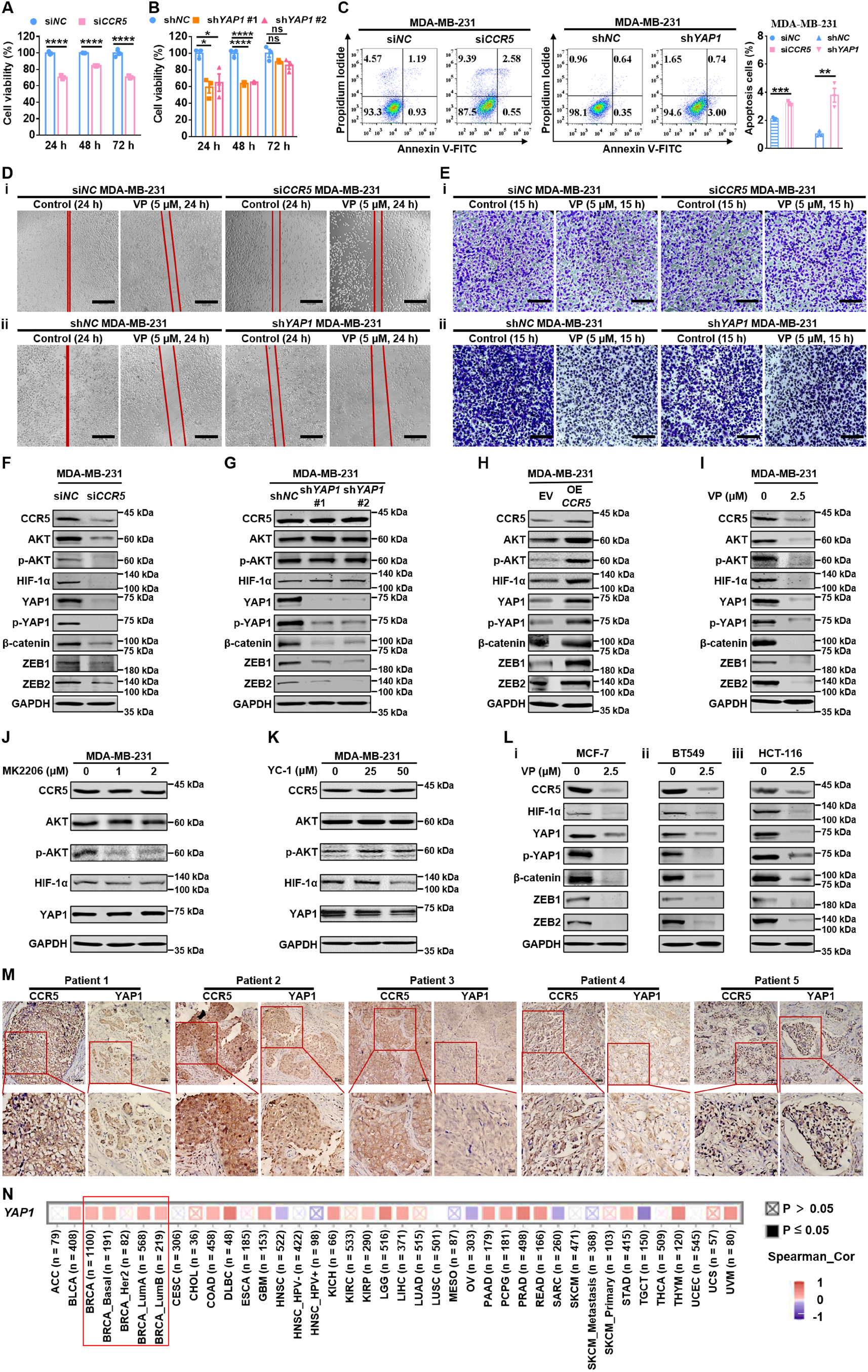
CCR5 is a key target of verteporfin and promotes metastasis via YAP1. (A-B) The cell viability was measured at the indicated time by MTS assay in control (si*NC*) and *CCR5* knockdown (si*CCR5*) (A), as well as control (sh*NC*) and *YAP1* knockdown (sh*YAP1* #1 and sh*YAP1* #2) MDA- MB-231 cells (B) (n = 3). (C) The apoptosis was analyzed by flow cytometry and quantified (n = 3) in si*NC* and si*CCR5*, as well as sh*NC* and sh*YAP1* MDA-MB-231 cells. (D) Representative images of scratch assay of the control (0.1% DMSO) and 5 μM verteporfin (VP) treatment in si*NC* and si*CCR5*, as well as sh*NC* and sh*YAP1* MDA-MB-231 cells. Scale bar: 520 μm. (E) Representative images of the transwell invasion assay of the control (0.1% DMSO) and 5 μM VP treatment in si*NC* and si*CCR5*, as well as sh*NC* and sh*YAP1* MDA-MB-231 cells. Scale bar: 220 μm. (F-I) The protein level of CCR5, AKT, phosphorylated-AKT (p-AKT), HIF-1α, YAP1, phosphorylated-YAP1 (p-YAP1), β-catenin, ZEB1 and ZEB2 was monitored by western blotting in si*NC* and si*CCR5* (F), sh*NC*, sh*YAP1* #1 and sh*YAP1* #2 (G), as well as EV (empty vector) and *CCR5* over expression (OE *CCR5*) MDA-MB-231 cells (H), and 48 h of VP (0 or 2.5 μM) treatment in MDA-MB-231 cells (I). (J-K) The protein level of CCR5, AKT, p-AKT, HIF-1α and YAP1 was monitored by western blotting in 48 h of AKT inhibitor MK2206 (0, 1 or 2 μM) treatment (J), or 48 h of HIF-1α inhibitor YC-1 (0, 25 or 50 μM) treatment in MDA-MB-231 cells (K). (L) The protein level of CCR5, HIF-1α, YAP1, p-YAP1, β-catenin, ZEB1 and ZEB2 was monitored by western blotting in 48 h of VP (0 or 2.5 μM) treatment in MCF-7 (i), BT549 (ii) and HCT-116 cells (iii). (M) Representative images of the co-expression of CCR5 and YAP1 in TNBC patient samples as analyzed by IHC. Scale bar: 50 μm for the 40 × vision and 20 μm for the zoomed vision. (N) The co-expression of *CCR5* and *YAP1* in different cancers were analyzed by the TIMER 2.0 website. Data in (A, and C) were analyzed by Unpaired two-tailed Student’s t test, and data in (B) was analyzed by one-way ANOVA. Data are presented as mean ± SEM. \**P* < 0.05, \*\**P* < 0.01, \*\*\**P* < 0.001, \*\*\*\**P* < 0.0001, ns: not significant.

Mechanistically, strongly reduced protein levels of epithelial-mesenchymal transition (EMT) markers β-catenin, Zinc finger E-box binding homeobox 1 (ZEB1) and Zinc finger E-box binding homeobox 2 (ZEB2) were observed in either *CCR5* or *YAP1* knockdown MDA-MB-231 cells (Fig. 7F and G, and Fig. S5C and D), and the reversed phenotype was detected in *CCR5*-overexpressing cells (Fig. 7H and Fig. S5E), suggesting that CCR5 positively regulated the expression of YAP1, which in turn regulated the expression of β-catenin, ZEB1 and ZEB2 in TNBC cells. Since it has been reported that CCR5 signaling can activate protein kinases such as AKT, as well as transcriptional regulators like hypoxia-inducible factor-1α (HIF-1α) [51], and HIF-1α can further directly bind to the YAP1 promoter in pancreatic ductal adenocarcinoma (PDAC) model [52], we then assessed these molecules in MDA-MB-231 cells. Unexpectedly, although the protein level of AKT, phosphorylated-AKT (p-AKT), HIF- 1α, YAP1, phosphorylated-YAP1 (p-YAP1) were all decreased in *CCR5* knockdown cells or with verteporfin treatment (Fig. 7F and I, and Fig. S5C and F), the protein level of YAP1 was not apparently changed when treated with AKT inhibitor MK2206 (Fig. 7J and Fig. S5G). However, it was decreased by the treatment with the HIF-1α inhibitor YC-1 (Fig. 7K and Fig. S5H), indicating that CCR5- dependent transcriptional regulation of YAP1 is not mediated by the AKT signaling pathway but via HIF-1α.

Interestingly, the addition of verteporfin reduced the expression of CCR5 in both mRNA and protein level (Fig. S5I and J), even for the induced CCR5 by CCL5 (Fig. S5K and L), however, the addition of another YAP1 inhibitor TED-347 didn’t affect the expression of CCR5 (Fig. S5J), hence, the reduced YAP1, p-YAP1, β-catenin, ZEB1 and ZEB2 proteins were observed with the treatment of verteporfin like *CCR5* knockdown in TNBC cells (Fig. 7F and I, and Fig. S5C and F), indicating that CCR5 is a new key target of verteporfin, which could regulate YAP1 via HIF-1α. Moreover, using verteporfin as a molecular probe, we further characterized the above signaling pathway proteins in BC cells of MCF- 7, TNBC cells of BT549 and colon cancer cells of HCT-116, the same regulation patten was observed in all these cancer cells (Fig. 7L and Fig. S5M-O), suggesting that CCR5-YAP1- ZEB1/ ZEB2/β- catenin signaling pathway is shared in BC, TNBC and colon cancer cells metastasis.

In order to further elucidate the co-expression of CCR5 and YAP1, IHC was performed in 10 TNBC patient samples. Notably, a positive correlation between the two proteins was noted (Fig. 7M). This result was also confirmed by the analysis of the co-expression of *CCR5* and *YAP1* in patients of different cancers as analyzed by the TIMER 2.0 website. The co-expression of *CCR5* and *YAP1* is presented in most of breast cancer patients, including BRCA, BRCA_Basal, BRCA_LumA and BRCA_LumB, except the BRCA_HER2 patients (Fig. 7N). Moreover, the co-expression is not limited to breast cancer; it could also be found in fifteen other cancers, such as thymoma (THYM), pancreatic ductal adenocarcinoma (PAAD), colon adenocarcinoma (COAD), low-grade glioma (LGG), and diffuse large B-cell lymphoma (DLBC). These data suggested that CCR5 regulated YAP1 expression might be presented in many cancer types.

## 4. Discussion

TNBC is a highly lethal cancer with few effective therapeutic strategies available at present [2], while drug resistance in BC is also increasing [53, 54]. CCL5/CCR5 axis plays an important role in cancer progression, including the tumor growth, drug resistance, and it has received considerable attention as a target for cancer therapy as well as anti-virus therapy [3-7, 55], including the intractable carcinomas such as TNBC [56] and pancreas cancer [57].

Maraviroc, the only FDA approved antagonist of CCR5 that can disrupt the interaction of CCR5 with HIV virus envelope glycoprotein gp120, has been widely used for functional studies of the CCL5/CCR5 axis with or without combination treatment [6, 58, 59]. Leronlimab (PRO140), a blocking antibody against CCR5 [60, 61], is currently used in combination with carboplatin for a clinical study of CCR5^+^ TNBC metastasis (NCT03838367). Both molecules are also involved in clinical studies for COVID-19 therapy (NCT04435522, NCT04710199, NCT04678830, NCT04347239 and NCT04343651). However, specific antagonists of the CCL5/CCR5 axis have not been reported, which has hampered efforts to elucidate the pathological functions and mechanisms of this axis in different diseases, as well as the development of therapeutic modalities against this target.

In this study, we developed an HTS assay for discovering the antagonists of the CCL5/CCR5 axis for the first time, which had a Z-factor of about 0.84, indicating it reached to the “excellent assay” [28]. Moreover, this assay can be performed using affordable reagents and standard laboratory instruments, making it accessible to a wide range of researchers. Additionally, by modifying the chemokines or their receptors, or utilizing fluorescence-labeled detection antibodies, the assay can be easily adapted to investigate regulators of other chemokine ligand/receptor axes, and further enhancing its versatility and sensitivity.

The strategy of drug repositioning or repurposing, which involves repurposing existing drugs for new therapeutic indications, has gained significant attention as a cost-effective and time-efficient alternative to *de novo* drug development [62, 63]. By leveraging our developed HTS assay and screening 7,555 compounds, including FDA-approved and NCI sets, we identified verteporfin as a novel antagonist of the CCL5/CCR5 axis. Verteporfin, originally approved by the FDA for PDT targeting subfoveal choroidal neovascularization in age-related macular degeneration [64], exhibited remarkable efficiency and specificity as a *bona fide* antagonist of the CCL5/CCR5 axis without PDT. Compared to maraviroc, verteporfin exhibited over 200-fold increased potency in disrupting the binding of CCL5 to CCR5. This superior performance and specific targeting of the CCL5/CCR5 axis were further confirmed through evaluation against seven other related chemokine ligand/receptor axes and two other YAP1 inhibitors. To note, for both HTS and IC50 analyses, we used the stable CCR5- expressing HEK-293T cells, not the membrane preparation, and the binding signal of CCL5 to CCR5 was further amplified by the HRP-conjugated anti-His antibody. Hence, in this assay, we analyzed the binding of CCL5 to CCR5 expressed on cells directly, which maintains the correct folding of CCR5 better than the cell lysis method for preparing the membrane protein CCR5, and this difference may account for the different IC50 values for maraviroc [65]. In addition to verteporfin, merbromin was also identified in this screening as having the ability to reduce the binding of CCL5 to CCR5. However, this compound has well known severe and lethal toxicities [66, 67], hence, it was used only as a positive control for the HTS assay.

Furthermore, we present clear evidence that verteporfin without PDT could also work as an immunotherapeutic agent like antibodies to reduce tumor growth *in vivo* with multiple immune competent or deficient mouse models. Verteporfin markedly reduced TNBC tumor growth via cell extrinsic effects on the immune system in TME. It significantly increased the infiltration of lymphocytes, and markedly decreased the recruitment of immunosuppressive cells. The importance of NK and CD8^+^ T cells in verteporfin-induced tumor reductions were further corroborated by antibody-mediated depletion. The cytotoxic effects of verteporfin *in vivo* were observed by the decreased tumor proliferation and increased apoptosis, enhanced levels of anti-tumor cytokines, as well as decreased pro-tumor cytokines. In addition, verteporfin also significantly prolonged the overall survival of mice, and decreased the viability of different types of cancer cells, as well as the DDP-resistant MDA-MB- 231 cells. Notably, a previous study showed that 8 mg/kg/day maraviroc significantly decreased the 4T1 tumor growth [16], which is different from our results. The difference may be accounted for by the different number of inoculated tumor cells used in the two studies: 5 × 10^4^ cells in the publication versus 2 × 10^5^ cells in our study. It’s worth pointing out that several other studies have reported that maraviroc has little effects on primary tumor growth at this dose or even higher doses in various models [10, 68-70]. In contrast, 8 mg/kg/day verteporfin already have significant effects on reducing 4T1 tumor growth, demonstrating a superior therapeutic index compared to maraviroc.

Metastasis is a major clinical problem in human cancers [71]. Compared to maraviroc, verteporfin without PDT also significantly reduced both migration and invasion of human TNBC cells *in vitro*, and markedly inhibited lung metastasis *in vivo*. Previous studies have shown CCR5 inhibitors reduced the tumor metastasis in both immune deficient and immune competent mice [10, 72], which is consistent with our finding that verteporfin decreased TNBC metastasis *in vivo* by a tumor cell-intrinsic mechanism through the CCR5-HIF-1α-YAP1-ZEB1/ZEB2/β-catenin cascade, without a reliance on the immune system.

Given the evidence that a small molecule like verteporfin could overcome the resistance of heterogeneous tumor cells to antibody therapy, and its high efficiency in reducing both TNBC tumor growth and metastasis, prolonging survival, verteporfin and its analogs could provide a promising immunotherapeutic modality for TNBC without PDT treatment.

Although previously CCR5 and YAP1 have been reported to promote tumor growth, including breast cancer [11, 73], our work is the first to show that mechanistically, CCR5 is a new key target of verteporfin, and it is essential for the expression of the key hippo effector YAP1 oncogene in BC, TNBC, and colon cancer cells, as well as in TNBC patient samples. In clear cell renal cell carcinoma (ccRCC), a positive regulatory loop, SOX17^low^/YAP/TEAD1/CCL5/CCR5/STAT3, was identified in the ccRCC-TAM interaction [74]. However, we observed the co-expression of CCR5 and YAP1 in BRCA but not kidney renal clear cell carcinoma (KIRC), suggesting that the regulation between CCR5 and YAP1 maybe different between TNBC and ccRCC. Previously, verteporfin was known as an inhibitor of YAP1 [75]. Here we provide the first evidence that verteporfin, but not YAP1 inhibitors such as CA3 or TED-347, could efficiently disrupt the CCL5/CCR5 axis and inhibit the expression of CCR5, and further decrease the expression of YAP1 at both mRNA and protein levels via HIF-1α, but not AKT. YAP1 is known for its crucial role in the initiation of most solid tumors [73]. It is a key component in the hippo pathway, which is highly conserved across higher-order vertebrates that modulates key target genes to regulate a multitude of biological processes including cellular proliferation, survival, differentiation, cellular fate determination, organ size, and tissue homeostasis [76]. Here is also for the first time that we show a GPCR receptor CCR5 regulates the expression of YAP1 via HIF-1α.

Up to now, only few cell-intrinsic signaling pathways regulated by the CCL5/CCR5 axis in BC have been reported, including the PI3K/AKT and NF-κB pathways [5, 77]. Here, we have uncovered two new mechanisms whereby the CCL5/CCR5 axis regulated YAP1 expression for immune (cell-extrinsic) regulation of TNBC tumor growth and cell-intrinsic regulation of metastasis respectively. This tumor-centric axis influenced the immune system in an extrinsic manner potentially by regulating immune cell recruiting chemokines such as CXCL16 and CXCL8, which were revealed by RNA-seq with or without siRNA targeting CCR5 in MDA-MB-231, and the key EMT players (ZEB1, ZEB2, and β- catenin) for metastasis (Fig. 8). Compared to a previous RNA-seq analysis with CCR5^+^ and CCR5^-^ SUM-159 cells isolated by FACS sorting [5], more different enriched KEGG pathways were revealed in our study with MBA-MD-231 cells subjected to siRNA-mediated CCR5 silencing. The difference could be due to the different properties of the cell lines used in the two studies, e.g., MDA-MB-231 cells are more resistant to the chemotherapy of paclitaxel or 5-FU (5-fluorouracil) than SUM-159 [78]. Here, we show that CCR5 via YAP1 regulated the expression of chemokines including CXCL16 and CXCL8. Although tumor-produced CXCL16 has been reported to attract NK, CD4^+^ T, and CD8^+^ T cells to TME in colon and breast cancers [45-47], and CXCL8 helps recruiting immunosuppressive MDSCs, TAMS, and Tregs [48-50], it would be of interest and importance to definitively elucidate the role of these chemokines in CCR5-regulated immune cell recruitment as a rational next step.

**Fig. 8.**
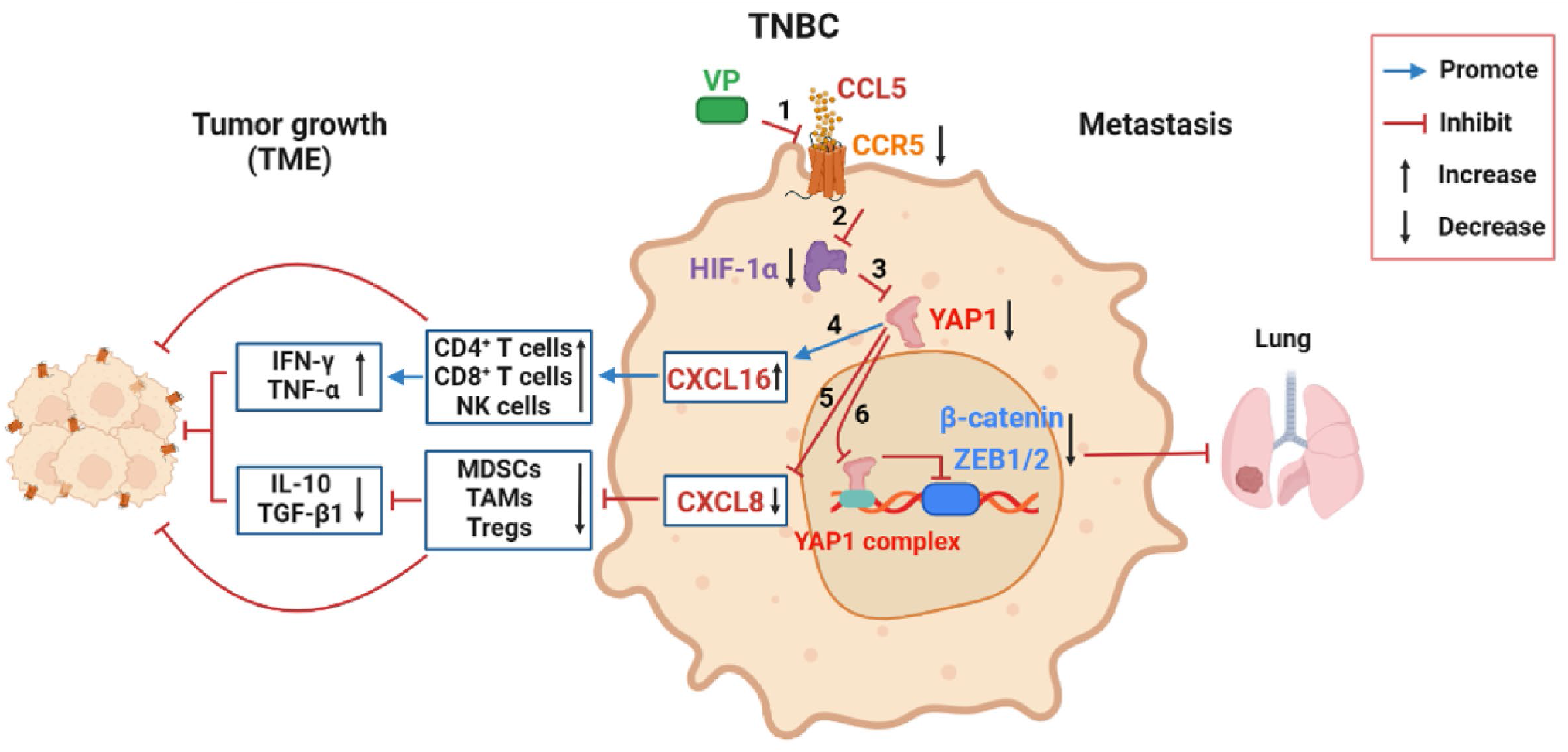
The mechanism of CCR5 regulated TNBC tumor growth in an immune dependent mechanism and metastasis in a cell-intrinsic way. In TNBC cells, the verteporfin (VP) treatment without PDT disrupted the binding of CCL5 to CCR5 and reduced the expression of CCR5 (1). The decreased CCR5 further led to the reduced HIF-1α (2), and decreased the expression of YAP1(3). The decreased YAP1 could up-regulate the expression of CXCL16 to further increase the recruitment of CD4^+^ T, CD8^+^ T and NK cells, as well as the anti-tumor cytokines like IFN-γ and TNF-α in TME (4). The decreased YAP1 could also down-regulate the expression of CXCL8 to further decrease the recruitment of immunosuppressive cells like MDSCs, TAMs and Tregs, as well as the pro-tumor cytokines like IL-10 and TGF-β1 in TME (5). With the regulation of both (4) and (5), VP, as well as CCR5, regulated the TNBC tumor growth mainly by immune regulation in TME. Moreover, the decreased YAP1 also inhibited the metastasis of TNBC to lung by down-regulating the expression of EMT markers β-catenin, ZEB1 and ZEB2 (6). Thus, VP targets CCL5/CCR5 axis to inhibit the TNBC growth by immune effects and metastasis by cell-intrinsic way via YAP1.

Using verteporfin as a molecular probe, we also observed the cell-intrinsic mechanism that CCR5 positively regulated YAP1 not only in BC, TNBC, and colon cancer cell lines, but also in TNBC patient samples. Furthermore, based on a TCGA database analysis, the co-expression of CCR5 and YAP1 is also found in basal, luminal A and luminal B subtypes of breast cancer except in HER2-positive breast cancer patients, as well as additional fifteen cancers, including THYM, PAAD, COAD, LGG, and DLBC, indicating that the CCL5/CCR5 mediated tumor growth and metastasis via YAP1 may operate in a wide range of malignant diseases.

In summary, our study provides for the first time a cell-based high-throughput screening (HTS) assay for identification of the antagonists targeting the CCL5/CCR5 axis, opening new avenues for studying other chemokine/receptor regulatory axes. Importantly, we present compelling evidence that verteporfin is a *bona fide* antagonist, it can serve as a molecular probe for the CCL5/CCR5 axis. Furthermore, we unveil a critical role of CCR5 in the regulation of YAP1 oncogene through HIF-1α and elucidate the immune regulatory mechanism of the CCL5/CCR5 axis in tumor growth via the CCR5-HIF-1α-YAP1-CXCL16/CXCL8 signaling cascade, as well as the cell-intrinsic pathway of CCR5-HIF-1α-YAP1-ZEB1/ZEB2/β-catenin in tumor metastasis. Notably, these regulatory mechanisms are likely to operate in various solid tumors. Additionally, we provide essential evidence that verteporfin, even without PDT, exhibits remarkable immunotherapeutic properties by significantly reducing tumor growth, inhibiting metastasis, and prolonging survival. These findings help deepen our understanding of the molecular mechanisms underlying the role of the CCL5/CCR5 axis in cancer progression and introduce a novel and promising immunotherapeutic agent for TNBC and potentially fifteen other malignancies characterized by high co-expression of CCR5 and YAP1 as well as inflammation, such as PAAD, COAD, LGG, and THYM.

## Ethical approval

The Institutional Animal Care and Use Committee of Shanghai Jiao Tong university (SJTU, Shanghai, China) approved this study.

## Consent for publication

All authors agree to the publication of the work entitled “A novel antagonist of the CCL5/CCR5 axis suppresses the tumor growth and metastasis of triple-negative breast cancer by CCR5-YAP1 regulation” to *Cancer Letters*.

## Availability of data and materials

The RNA-seq data from this publication have been deposited to the Gene Expression Omnibus (GEO) database and assigned the identifier GSE222052. It is publicly available as of the date of publication.

## Availability of supporting data

All data generated or analyzed during this study are included in this published article.

## CRediT authorship contribution statement

**Ling Chen:** Conceptualization, methodology, investigation, formal analysis, writing-original draft, writing-review & editing. **Guiying Xu:** Investigation, formal analysis, writing-review & editing. **Xiaoxu Song:** Investigation, writing-review & editing. **Lianbo Zhang:** Investigation, formal analysis, writing-review & editing. **Chuyu Chen:** Investigation, writing-review & editing. **Gang Xiang:** Investigation, writing-review & editing. **Shuxuan Wang:** Investigation, writing-review & editing. **Zijian Zhang:** Investigation, writing-review & editing. **Fang Wu:** Resources, writing-review & editing. **Xuanming Yang:** Resources, writing-review & editing. **Lei Zhang:** Resources, writing-review & editing. **Xiaojing Ma:** Conceptualization, methodology, resources, formal analysis, writing-original draft, writing-review & editing, supervision, funding acquisition. **Jing Yu:** Conceptualization, methodology, resources, formal analysis, writing-original draft, writing-review & editing, supervision, funding acquisition.

## Acknowledgments

We thank Jiahuai Han (Xiamen University, Xiamen, China) for providing the human *CCL5*, *CCL7*, *CCL11* and *CCR5* cDNA. We thank Haixia Jiang, Peide Cheng and Qian Luo from the Core Facility and Technical Service Center for SLSB, School of Life Sciences and Biotechnology, Shanghai Jiao Tong University for technical support with FACS and macroscopy analysis.

## Abbreviations

BC: Breast cancer
BRCA: Breast invasive carcinoma
COAD: Colon adenocarcinoma
COVID-19: Coronavirus disease 2019
DDP: Cisplatin
DEGs: Differentially expressed genes
DLBC: Diffuse large B-cell lymphoma
EMT: Epithelial-mesenchymal transition
ESCA: Esophageal carcinoma
FACS: Fluorescence-activated cell sorting
GBM: Glioblastoma multiforme
GEPIA: Gene expression profiling interactive analysis
GO: Gene ontology
HTS: High-throughput screening
KEGG: Kyoto Encyclopedia of Genes and Genomes
KIRC: Kidney renal clear cell carcinoma
LGG: Low-grade glioma
MDSCs: Myeloid-derived suppressor cells
NK: Natural killer
PAAD: Pancreatic adenocarcinoma
PDAC: Pancreatic ductal adenocarcinoma
PDT: Photodynamic treatment
shRNAs: Short hairpin RNAs
siRNA: Small interfering RNA
TAMs: Tumor-associated macrophages
THYM: Thymoma
TME: Tumor microenvironment
TNBC: Triple-negative breast cancer
Tregs: Regulatory T cells

