## Supplemental material for "A novel antagonist of the CCL5/CCR5 axis suppresses the tumor growth and metastasis of triple-negative breast cancer by CCR5-YAP1 regulation"

**Supplementary Table S1. Primers used for amplification of human genes**

| Primers | Sequences |
|---|---|
| *CCL3*-Forward | 5'-CGGAATTCCGATGCAGGTCTCCACTGCT-3' |
| *CCL3*-Reverse | 5'-CGGGATCCCGCTAATGGTGATGGTGATGATGGGGCCCCTGGAACAGAACTTCCAGGGCACTCAGCTCCAGG-3' |
| *CCL4*-Forward | 5'-CGGAATTCCGATGAAGCTCTGCGTGACTGT-3' |
| *CCL4*-Reverse | 5'-CGGGATCCCGCTAATGGTGATGGTGATGATGGGGCCCCTGGAACAGAACTTCCAGGTTCAGTTCCAGGTCATACACG-3' |
| *CCL5*-Forward | 5'-CGGAATTCCGATGAAGGTCTCCGCGGCA-3' |
| *CCL5*-Reverse | 5'-CGGGATCCCGCTAATGGTGATGGTGATGATGGGGCCCCTGGAACAGAACTTCCAGGCTCATCTCCAAAGAGTTGA-3' |
| *CCL7*-Forward | 5'-CGGAATTCCGATGAAAGCCTCTGCAGCA-3' |
| *CCL7*-Reverse | 5'-CGGGATCCCGTCAATGGTGATGGTGATGATGGGGCCCCTGGAACAGAACTTCCAGAAGCTTTGGAGTTTGG-3' |
| *CCL11*-Forward | 5'-CGGAATTCCGATGAAGGTCTCCGCAGC-3' |
| *CCL11*-Reverse | 5'-CGGGATCCCGCTAATGGTGATGGTGATGATGGGGCCCCTGGAACAGAACTTCCAGAGGCTTTGGAGTTGGAGA-3' |
| *CCR1*-Forward | 5'-GCTCTAGAGCATGGAAACTCCAAACACCACAGAG-3' |
| *CCR1*-Reverse | 5'-GGAATTCCCTACTTATCGTCGTCATCCTTGTAATCGGGCCCCTGGAACAGAACTTCCAGGAACCCAGCAGAGAGTTCATG-3' |
| *CCR3*-Forward | 5'-GCTCTAGAGCATGACAACCTCACTAGATACAGTTGA-3' |
| *CCR3*-Reverse | 5'-GGAATTCCCTACTTATCGTCGTCATCCTTGTAATCGGGCCCCTGGAACAGAACTTCCAGAAACACAATAGAGAGTTCCGG-3' |
| *CCR5*-Forward | 5'-GCTCTAGAGCATGGATTATCAAGTGTCAAGTC-3' |
| *CCR5*-Reverse | 5'-GGAATTCCCTACTTATCGTCGTCATCCTTGTAATCGGGCCCCTGGAACAGAACTTCCAGCAAGCCCACAGATATTTCCTG-3' |
| *GPR75*-Forward | 5'-GCTCTAGAGCATGAACTCAACAGGCCACC-3' |
| *GPR75*-Reverse | 5'-GGAATTCCTTACTTATCGTCGTCATCCTTGTAATCGGGCCCCTGGAACAGAACTTCCAGAACGGAGGGGACTGGAA-3' |

**Supplementary Table S2. Sequences of siRNAs and shRNAs**

| **Primers** | **Sequences** |
|---|---|
| Non-target siRNA (sense) | 5’-UUCUCCGAACGUGUCACGUTT-3’ |
| Human *CCR5* siRNA (sense) | 5’-GGGAGAAGUUCAGAAACUATT-3’ |
| Scrambled shRNA (sense) | 5’-CAACAAGATGAAGAGCACCAA-3’ |
| Human *YAP1* shRNA1 (sense) | 5’-CCCAGTTAAATGTTCACCAAT-3’ |
| Human *YAP1* shRNA2 (sense) | 5’-GCCACCAAGCTAGATAAAGAA-3’ |

**Supplementary Table S3. Primers used for qPCR**

| **Primers** | **Sequences** |
|---|---|
| Human *CCR5*-Forward | 5'-ACTGCAAAAGGCTGAAGAGC-3' |
| Human *CCR5*-Reverse | 5'-AGCATAGTGAGCCCAGAAGG-3' |
| Human *YAP1*-Forward | 5'-TAGCCCTGCGTAGCCAGTTA-3' |
| Human *YAP1*-Reverse | 5'-TCATGCTTAGTCCACTGTCTGT-3' |
| Human *CXCL8*-Forward | 5'-GAGAGTGATTGAGAGTGGACCAC-3' |
| Human *CXCL8*-Reverse | 5'-CACAACCCTCTGCACCCAGTTT-3' |
| Human *CXCL16*-Forward | 5'-CCTATGTGCTGTGCAAGAGGAG-3' |
| Human *CXCL16*-Reverse | 5'-CTGGGCAACATAGAGTCCGTCT-3' |
| Human *GAPDH*-Forward | 5'-GGAGCGAGATCCCTCCAAAAT-3' |
| Human *GAPDH*-Reverse | 5'-GGCTGTTGTCATACTTCTCATGG-3' |

**Supplementary Table S4. Antibodies used in the study**

| Antibody | Dilution | Vendor | Catalog | RRID |
|---|---|---|---|---|
| HIF-1α | 1:1000 | Abbkine | ABP51513 | AB_2927791 |
| Integrin alpha 2 | 1:500 | Abcam | ab181548 | AB_2847852 |
| p-YAP1 (S127) | 1:1000 | Abmart | T55743 | AB_2927792 |
| CXCL8 (IL8) | 1:250 | Abmart | TD6998 | AB_2934038 |
| CCR5 | 1:1000 | Affinity Biosciences | AF6339 | AB_2835195 |
| CXCL16 | 1:500 | Affinity Biosciences | DF13312 | AB_2846331 |
| β-Catenin | 1:1000 | BD Biosciences | 610153 | AB_397554 |
| Alexa Fluor 488-labeled Goat Anti-Mouse IgG (H+L) | 1:500 | Beyotime Biotechnology | A0428 | AB_2893435 |
| CD16/32 | 1:300 | Biolegend | 101335 | AB_2563723 |
| Akt (pan) | 1:1000 | Cell Signaling Technology | 4685 | AB_2225340 |
| p-Akt (Ser473) | 1:1000 | Cell Signaling Technology | 4060 | AB_2315049 |
| IRDye® 800CW Goat anti-Mouse IgG | 1:5000 | LI-COR Biosciences | 926-32210 | AB_621842 |
| IRDye® 800CW Donkey anti-Rabbit IgG | 1:5000 | LI-COR Biosciences | 926-32213 | AB_621848 |
| HRP-conjugated 6*His, His-Tag | 1:5000 | Proteintech | HRP-66005 | AB_2857904 |
| 6*His, His-Tag | 1:1000 | Proteintech | 66005-1-Ig | AB_11232599 |
| YAP1 | 1:1000 | Proteintech | 66900-1-Ig | AB_2882229 |
| ZEB1 | 1:1000 | Proteintech | 21544-1-AP | AB_10734325 |
| ZEB2 | 1:1000 | Proteintech | 14026-1-AP | AB_2878001 |
| GAPDH | 1:1000 | Santa Cruz Biotechnology | sc-47724 | AB_627678 |
| Ki67 | 1:500 | Servicebio | GB111499 | AB_2927572 |
| Ly6G | 1:300 | Servicebio | GB11229 | AB_2814689 |
| Foxp3 | 1:100 | Servicebio | GB11093 | AB_2861434 |
| CD4 | 1:200 | Servicebio | GB13064-2 | AB_2892096 |

| | | | | |
|---|---|---|---|---|
| CD8 | 1:300 | Servicebio | GB13429 | AB_2904188 |
| HRP conjugated Goat Anti-Rabbit IgG (H+L) | 1:200 | Servicebio | GB23303 | AB_2811189 |
| *In Vivo* Plus anti-mouse CD8α | N/A | BioXCell | BP0061 | AB_1125541 |
| Anti-Mouse Asialo GM1 | N/A | CEDARLANE | CL8955 | AB_10090118 |
| PE Hamster Anti-Mouse CD11c | 1:500 | BD Biosciences | 557401 | AB_396684 |
| FITC Rat Anti-Mouse I-A/I-E | 1:500 | BD Biosciences | 562009 | AB_10893593 |
| FITC anti-mouse/human CD11b | 1:500 | Biolegend | 101205 | AB_312788 |
| PE anti-mouse Ly-6G/Ly-6C (Gr-1) | 1:500 | Biolegend | 108407 | AB_313372 |
| PerCP/Cyanine5.5 anti-mouse Ly-6C | 1:500 | Biolegend | 128011 | AB_1659242 |
| APC anti-mouse Ly-6G | 1:500 | Biolegend | 127613 | AB_1877163 |
| PerCP/Cyanine5.5 anti-mouse CD45 | 1:500 | Biolegend | 103131 | AB_893344 |
| FITC anti-mouse CD45 | 1:500 | Biolegend | 157214 | AB_2894427 |
| PE anti-mouse CD4 | 1:500 | Biolegend | 100408 | AB_312693 |
| PerCP/Cyanine5.5 anti-mouse CD4 | 1:500 | Biolegend | 100434 | AB_893324 |
| PE anti-mouse CD8a | 1:500 | Biolegend | 100707 | AB_312746 |
| PE anti-mouse CD49b | 1:500 | Biolegend | 108907 | AB_313414 |
| APC anti-mouse CD3 | 1:500 | Biolegend | 100236 | AB_2561456 |
| APC anti-mouse CD106 | 1:500 | Biolegend | 105717 | AB_1877142 |
| FITC anti-human CD195 (CCR5) | 1:500 | Biolegend | 313705 | AB_345305 |
| PE anti-mouse CD25 | 1:500 | Biolegend | 102007 | AB_312856 |
| Alexa Fluor® 488 anti-mouse/rat/human FOXP3 | 1:500 | Biolegend | 320011 | AB_439747 |
| PerCP/Cyanine5.5 anti-mouse F4/80 | 1:500 | Biolegend | 123127 | AB_893496 |
| Alexa Fluor® 647 anti-mouse CD206 (MMR) | 1:500 | Biolegend | 141712 | AB_10900420 |

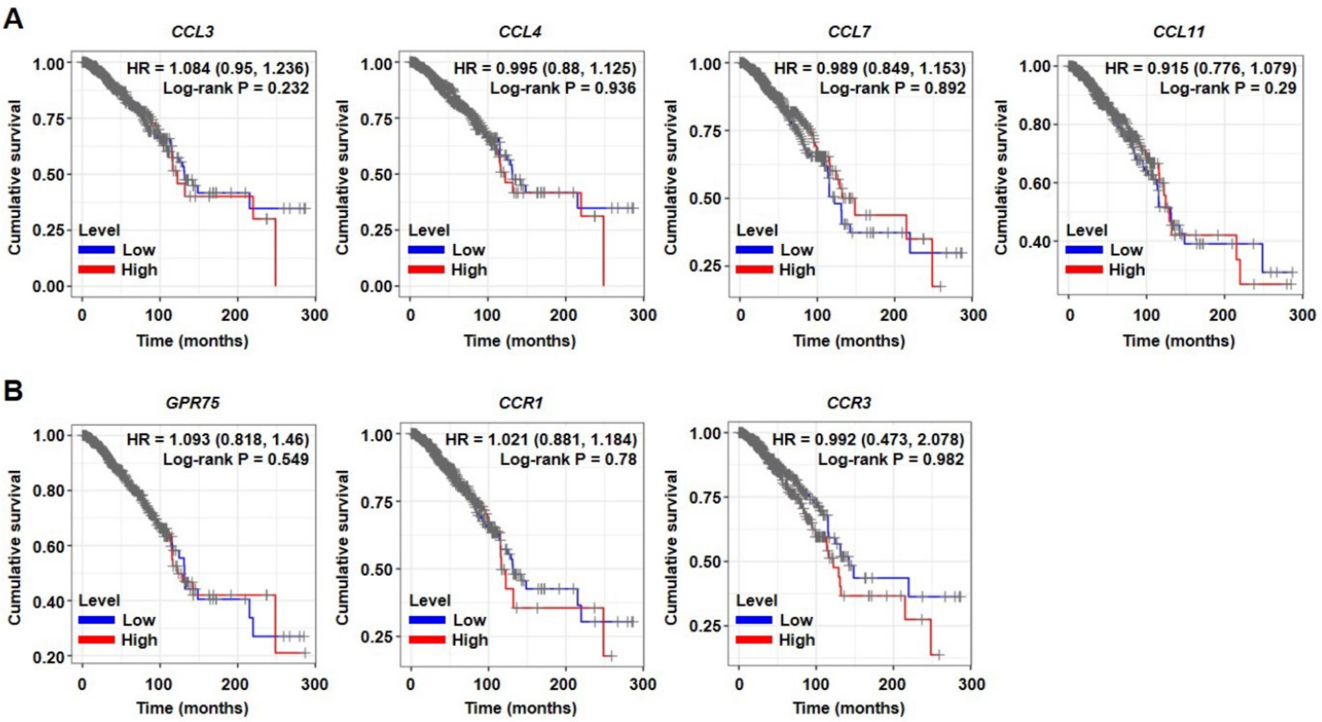


**Fig. S1. The high and low expression of the other ligands for CCR5 (A) and the other receptors for CCL5 (B) are not associated with BRCA patients' cumulative survival as analyzed by the TIMER website, respectively.**

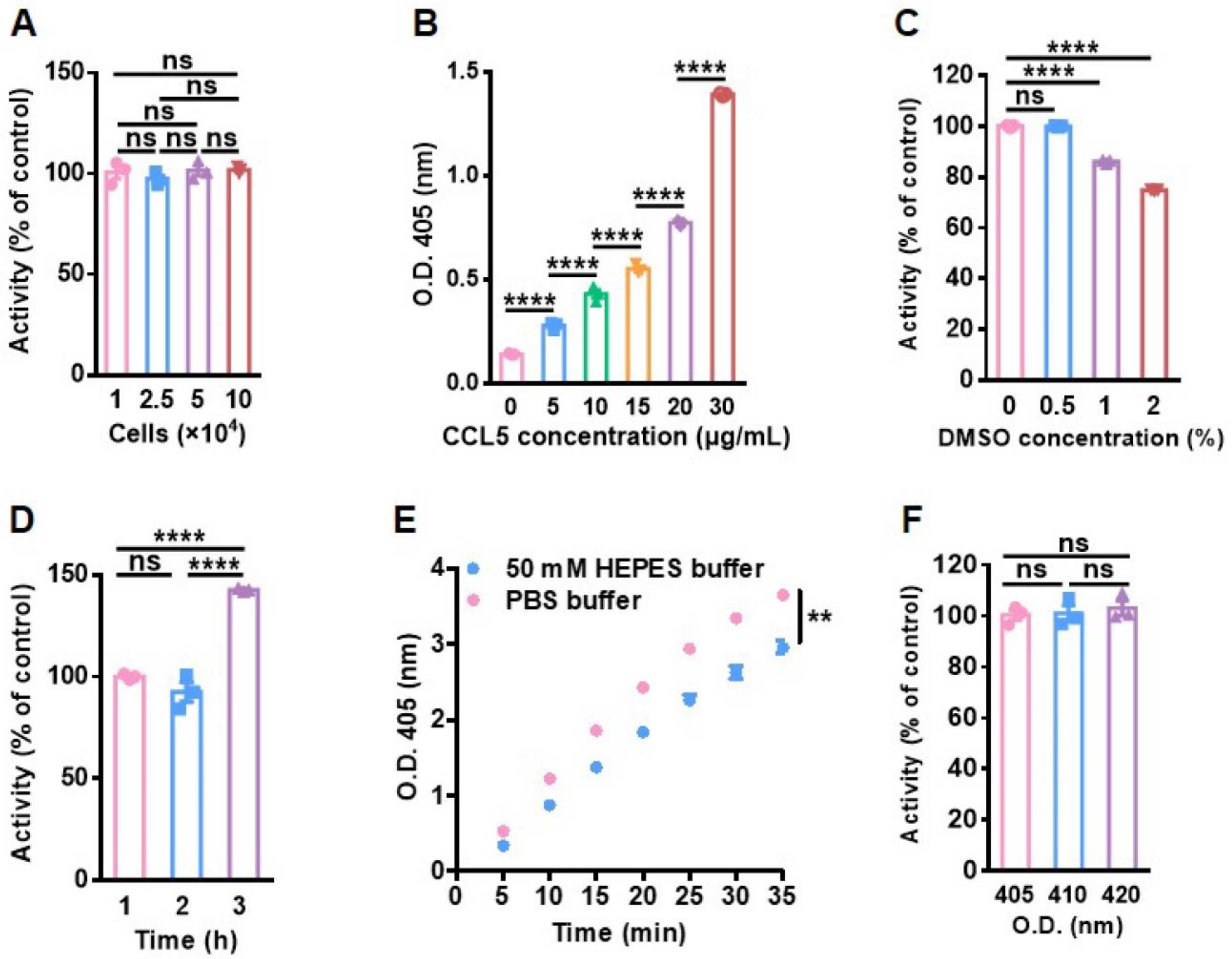


**Fig. S2. The optimization of the high-throughput screening (HTS) assay for discovering antagonists of CCL5/CCR5 axis.** (A-F) The numbers of CCR5 stable expressing cells (A), the concentration of His-tagged CCL5 peptides (B), as well as DMSO (C), the binding time of CCL5 with CCR5 (D), the assay buffer (E) and the detecting wave (F) were optimized for developing the HTS assay (n = 3). Data in (A-D, and F) were analyzed by one-way ANOVA, and data in (E) was analyzed by unpaired two-tailed Student's t test. Data were reported as means ± SEM. ***P* < 0.01, *****P* < 0.0001, ns: not significant.

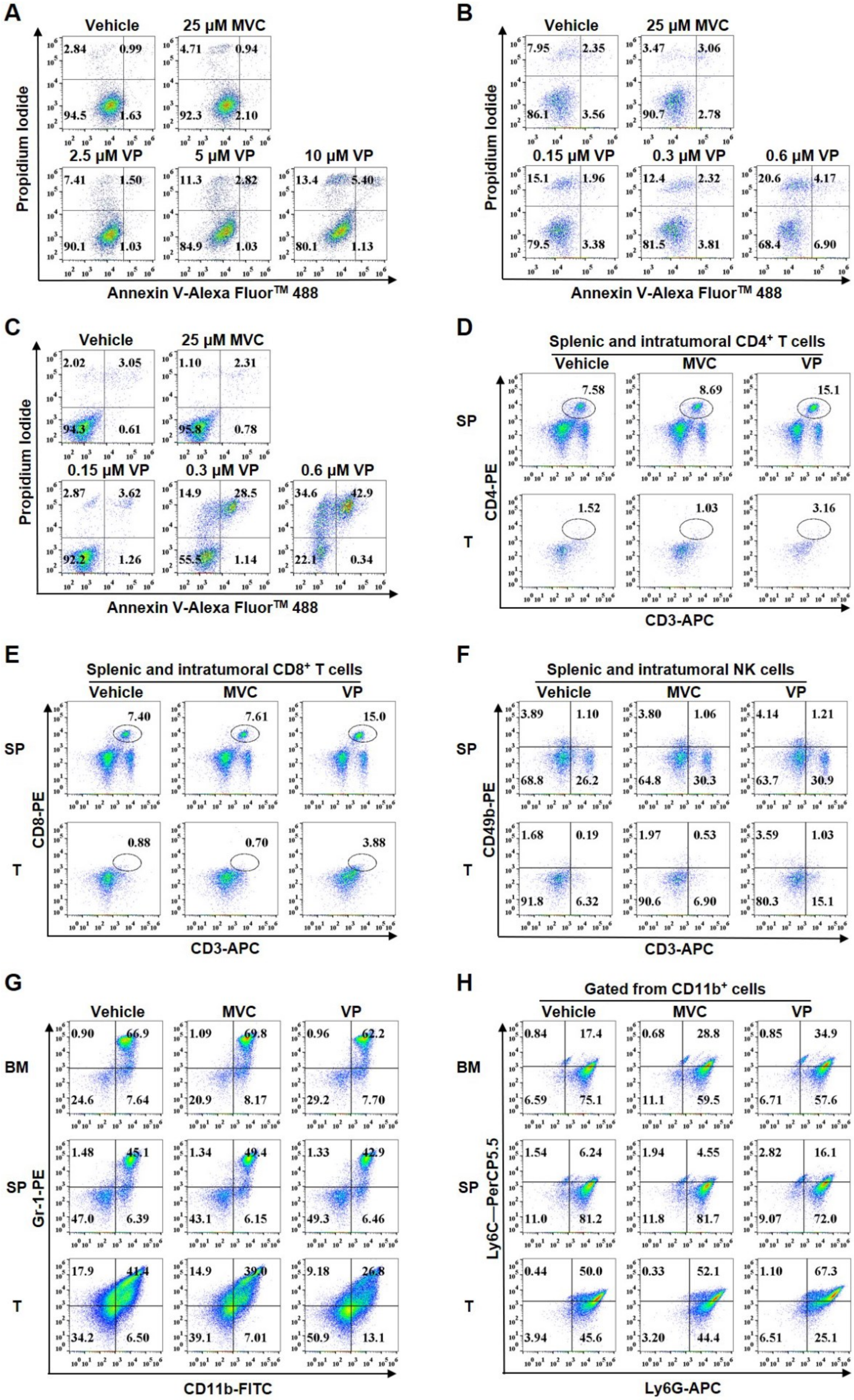

A
Vehicle
25 μM MVC
2.5 μM VP
5 μM VP
10 μM VP
2.84 0.99 94.5 1.63
4.71 0.94 92.3 2.10
7.41 1.50 90.1 1.03
11.3 2.82 84.9 1.03
13.4 5.40 80.1 1.13
Propidium Iodide
Annexin V-Alexa Fluor™ 488
B
Vehicle
25 μM MVC
0.15 μM VP
0.3 μM VP
0.6 μM VP
7.95 2.35 86.1 3.56
3.47 3.06 90.7 2.78
15.1 1.96 79.5 3.38
12.4 2.32 81.5 3.81
20.6 4.17 68.4 6.90
Propidium Iodide
Annexin V-Alexa Fluor™ 488
C
Vehicle
25 μM MVC
0.15 μM VP
0.3 μM VP
0.6 μM VP
2.02 3.05 94.3 0.61
1.10 2.31 95.8 0.78
2.87 3.62 92.2 1.26
14.9 28.5 55.5 1.14
34.6 42.9 22.1 0.34
Propidium Iodide
Annexin V-Alexa Fluor™ 488
D
Splenic and intratumoral CD4+ T cells
Vehicle
MVC
VP
SP
7.58 8.69 15.1
T
1.52 1.03 3.16
CD4-PE
CD3-APC
E
Splenic and intratumoral CD8+ T cells
Vehicle
MVC
VP
SP
7.40 7.61 15.0
T
0.88 0.70 3.88
CD8-PE
CD3-APC
F
Splenic and intratumoral NK cells
Vehicle
MVC
VP
SP
3.89 1.10 68.8 26.2
3.80 1.06 64.8 30.3
4.14 1.21 63.7 30.9
T
1.68 0.19 91.8 6.32
1.97 0.53 90.6 6.90
3.59 1.03 80.3 15.1
CD49b-PE
CD3-APC
G
Vehicle
MVC
VP
BM
0.90 66.9 24.6 7.64
1.09 69.8 20.9 8.17
0.96 62.2 29.2 7.70
SP
1.48 45.1 47.0 6.39
1.34 49.4 43.1 6.15
1.33 42.9 49.3 6.46
T
17.9 41.4 34.2 6.50
14.9 39.0 39.1 7.01
9.18 26.8 50.9 13.1
Gr-1-PE
CD11b-FITC
H
Gated from CD11b+ cells
Vehicle
MVC
VP
BM
0.84 17.4 6.59 75.1
0.68 28.8 11.1 59.5
0.85 34.9 6.71 57.6
SP
1.54 6.24 11.0 81.2
1.94 4.55 11.8 81.7
2.82 16.1 9.07 72.0
T
0.44 50.0 3.94 45.6
0.33 52.1 3.20 44.4
1.10 67.3 6.51 25.1
Ly6C—PerCP5.5
Ly6G-APC

**Fig. S3. The analysis of cell apoptosis in vitro and the immune cells in vivo post maraviroc (MVC) or verteporfin (VP) treatment without PDT by flow cytometry.** (A-C) The apoptosis of MDA-MB-231 (A), BT549 (B) and 4T1 cells (C) treated with MVC or VP at the indicated concentration was analyzed by flow cytometry, respectively. (D-F) The $CD4^+$ T (D), $CD8^+$ T (E) and NK cells (F) in spleens (SP) and tumors (T) were analyzed on day 14 post tumor injection in the 4T1 tumor-bearing mice treated with vehicle, MVC or VP, respectively. (G-H) Total MDSCs (G) and Ly6G/Ly6C expression cells (H) in bone marrow (BM), SP and T were analyzed on day 28 post tumor inoculation in 4T1 tumor-bearing mice treated with vehicle, MVC or VP, respectively.

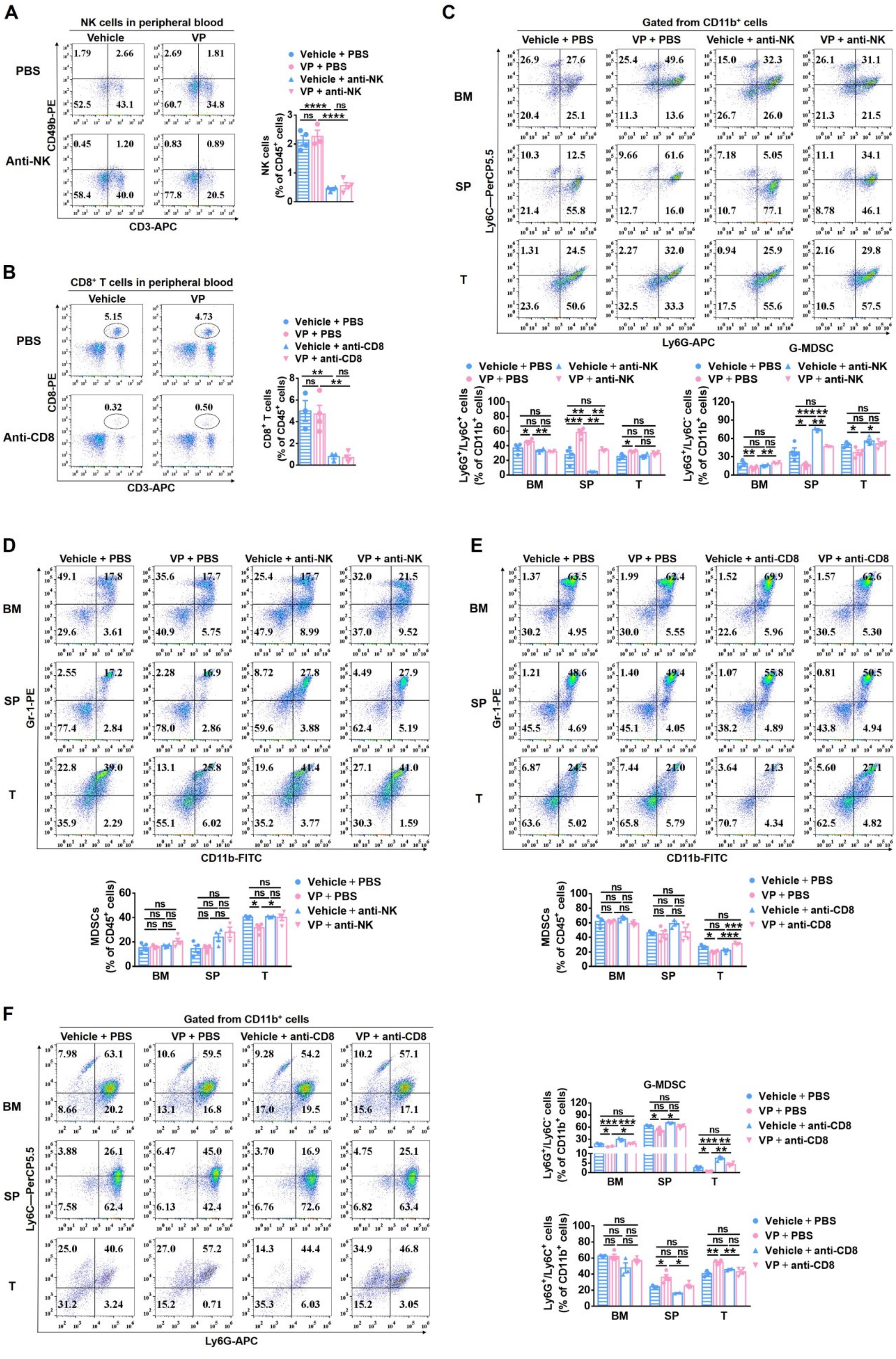

A
NK cells in peripheral blood
Vehicle
VP
PBS
Anti-NK
CD49b-PE
CD3-APC
1.79
2.66
52.5
43.1
2.69
1.81
60.7
34.8
0.45
1.20
58.4
40.0
0.83
0.89
77.8
20.5
Vehicle + PBS
VP + PBS
Vehicle + anti-NK
VP + anti-NK
NK cells (% of CD45+ cells)
B
CD8+ T cells in peripheral blood
Vehicle
VP
PBS
Anti-CD8
CD8-PE
CD3-APC
5.15
4.73
0.32
0.50
Vehicle + PBS
VP + PBS
Vehicle + anti-CD8
VP + anti-CD8
CD8+ T cells (% of CD45+ cells)
C
Gated from CD11b+ cells
Vehicle + PBS
VP + PBS
Vehicle + anti-NK
VP + anti-NK
BM
SP
T
Ly6C—PerCP5.5
Ly6G-APC
26.9
27.6
20.4
25.1
25.4
49.6
11.3
13.6
15.0
32.3
26.7
26.0
26.1
31.1
21.3
21.5
10.3
12.5
21.4
55.8
9.66
61.6
12.7
16.0
7.18
5.05
10.7
77.1
11.1
34.1
8.78
46.1
1.31
24.5
23.6
50.6
2.27
32.0
32.5
33.3
0.94
25.9
17.5
55.6
2.16
29.8
10.5
57.5
G-MDSC
Ly6G+/Ly6C+ cells (% of CD11b+ cells)
Ly6G+/Ly6C- cells (% of CD11b+ cells)
D
Vehicle + PBS
VP + PBS
Vehicle + anti-NK
VP + anti-NK
BM
SP
T
Gr-1-PE
CD11b-FITC
49.1
17.8
29.6
3.61
35.6
17.7
40.9
5.75
25.4
17.7
47.9
8.99
32.0
21.5
37.0
9.52
2.55
17.2
77.4
2.84
2.28
16.9
78.0
2.86
8.72
27.8
59.6
3.88
4.49
27.9
62.4
5.19
22.8
39.0
35.9
2.29
13.1
25.8
55.1
6.02
19.6
41.4
35.2
3.77
27.1
41.0
30.3
1.59
MDSCs (% of CD45+ cells)
E
Vehicle + PBS
VP + PBS
Vehicle + anti-CD8
VP + anti-CD8
BM
SP
T
Gr-1-PE
CD11b-FITC
1.37
63.5
30.2
4.95
1.99
62.4
30.0
5.55
1.52
69.9
22.6
5.96
1.57
62.6
30.5
5.30
1.21
48.6
45.5
4.69
1.40
49.4
45.1
4.05
1.07
55.8
38.2
4.89
0.81
50.5
43.8
4.94
6.87
24.5
63.6
5.02
7.44
21.0
65.8
5.79
3.64
21.3
70.7
4.34
5.60
27.1
62.5
4.82
MDSCs (% of CD45+ cells)
F
Gated from CD11b+ cells
Vehicle + PBS
VP + PBS
Vehicle + anti-CD8
VP + anti-CD8
BM
SP
T
Ly6C—PerCP5.5
Ly6G-APC
7.98
63.1
8.66
20.2
10.6
59.5
13.1
16.8
9.28
54.2
17.0
19.5
10.2
57.1
15.6
17.1
3.88
26.1
7.58
62.4
6.47
45.0
6.13
42.4
3.70
16.9
6.76
72.6
4.75
25.1
6.82
63.4
25.0
40.6
31.2
3.24
27.0
57.2
15.2
0.71
14.3
44.4
35.3
6.03
34.9
46.8
15.2
3.05
G-MDSC
Ly6G+/Ly6C- cells (% of CD11b+ cells)
Ly6G+/Ly6C+ cells (% of CD11b+ cells)

**Fig. S4. The analysis of NK or CD8$^+$ T cells' depletion efficiency in peripheral blood and the MDSCs in vivo post verteporfin (VP) treatment without PDT by flow cytometry.** (A-B) The NK cells (A) or CD8$^+$ T cells (B) in peripheral blood were analyzed and quantified (n = 4) in 4T1 tumor-bearing BALB/c mice treated with or without anti-mouse/rat asialo GM1 (anti-NK) antibody, or anti-CD8α (anti-CD8) antibody, respectively. (C-D) Ly6G/Ly6C expression cells (C) and total MDSCs (D) in bone marrow (BM), spleens (SP) and tumors (T) were analyzed on day 14 post tumor inoculation in 4T1 tumor-bearing mice treated with or without anti-NK antibody, or combined with vehicle or VP, respectively (n = 4). (E-F) Total MDSCs (E) and Ly6G/Ly6C expression cells (F) in BM, SP and T were analyzed on day 22 post tumor inoculation in 4T1 tumor-bearing mice treated with or without anti-CD8 antibody, or combined with vehicle or VP, respectively (n = 4). Data in (A-F) were analyzed by One-way ANOVA. Data are presented as mean ± SEM. *$P < 0.05$, **$P < 0.01$, ***$P < 0.001$, ****$P < 0.0001$, ns: not significant.

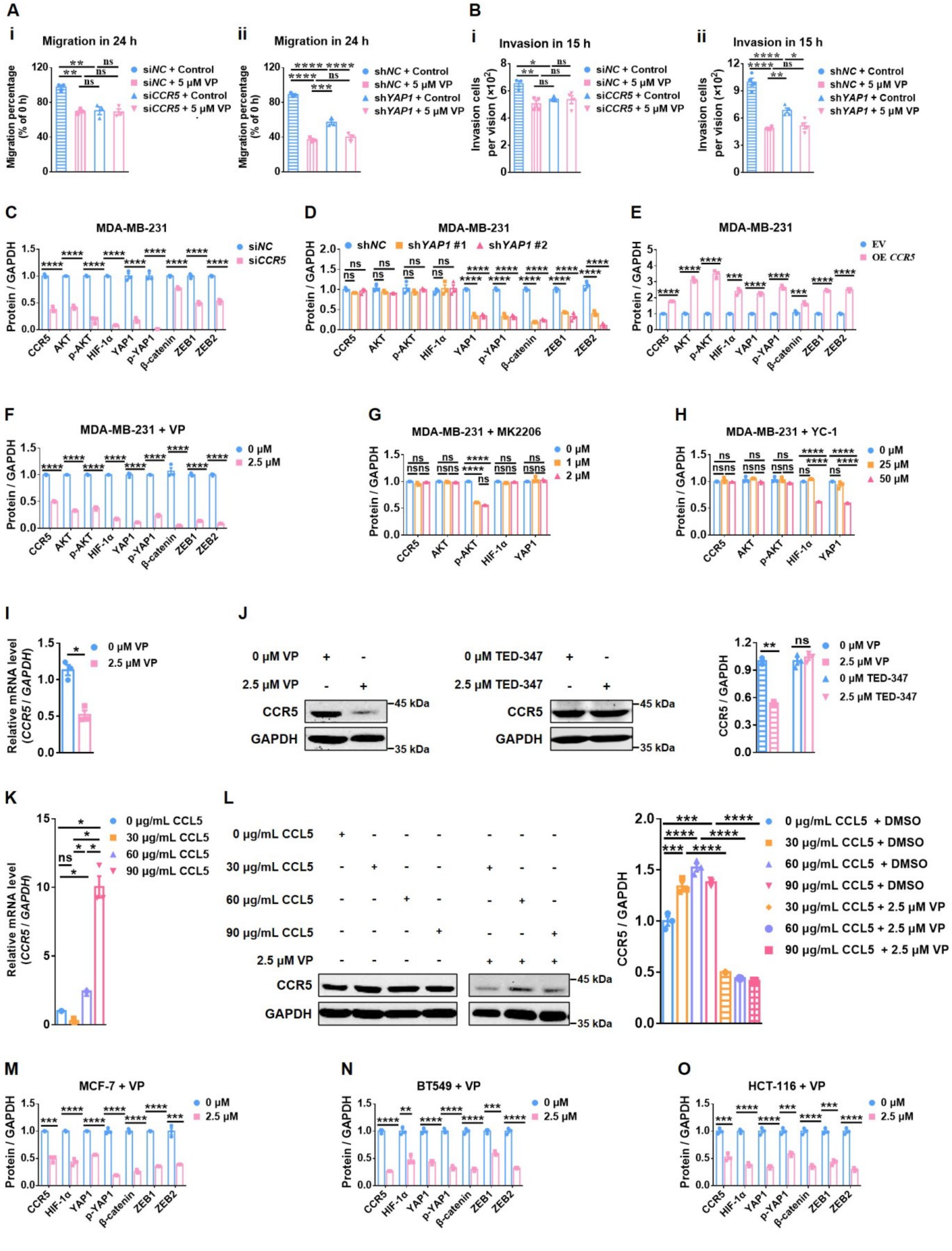


**Fig. S5. CCR5 is a key target of verteporfin and promotes metastasis via YAP1.** (A-B)

Quantification of the migration percentage (n = 3) (A) and invaded cells (n = 4) (B) under indicated treatment in control (si*NC*) and *CCR5* knockdown (si*CCR5*), as well as control (sh*NC*) and *YAP1* knockdown (sh*YAP1*) MDA-MB-231 cells in Figures 7D and 7E. (C-F) Quantification of the protein level of CCR5, AKT, p-AKT, HIF-1α, YAP1, p-YAP1, β-catenin, ZEB1 and ZEB2 monitored by western blotting in si*NC* and si*CCR5* (C), sh*NC* and *YAP1* knockdown (sh*YAP1* #1 and sh*YAP1* #2) (D), EV (empty vector) and *CCR5* over expression (OE *CCR5*) MDA-MB-231 cells (E), and 48 h of verteporfin (VP; 0 or 2.5 μM) treated MDA-MB-231 cells (F) in Figures 7F-7I (n = 3). (G-H) Quantification of the protein level of CCR5, AKT, p-AKT, HIF-1α and YAP1 monitored by western blotting in MK2206 (0, 1 or 2 μM) treated (G) or YC-1 (0, 25 or 50 μM) treated (H) MDA-MB-231 cells in Figures 7J and 7K (n = 3). (I) The mRNA expression of *CCR5* was measured by qPCR in VP (0 or 2.5 μM) treated MDA-MB-231 cells (n = 3). (J) The protein level of CCR5 was analyzed by western blotting and quantified (n = 3) in VP (0 or 2.5 μM) or TED-347 (0 or 2.5 μM) treated MDA-MB-231 cells. (K) The mRNA expression of *CCR5* was measured by qPCR in 72 h of CCL5 (0, 30, 60 or 90 μg/ml) treatment in MDA-MB-231 cells (n = 3). (L) The protein level of CCR5 was detected by western blotting and quantified (n = 3) in 72 h of CCL5 (0, 30, 60 or 90 μg/ml) treatment or CCL5 combined VP (0 or 2.5 μM) treatment in MDA-MB-231 cells. (M-O) Quantification of the protein level of CCR5, HIF-1α, YAP1, p-YAP1, β-catenin, ZEB1 and ZEB2 monitored by western blotting in 48 h of VP (0 or 2.5 μM) treatment in MCF-7 (M), BT549 (N) and HCT-116 cells (O) in Figure 7L (n = 3). Data in (C, E, F, I, J, and M-O) were analyzed by Unpaired two-tailed Student's t test, and data in (A, B, D, G, H, K, and L) were analyzed by One-way ANOVA. Data are presented as mean ± SEM. *$P < 0.05$, **$P < 0.01$, ***$P < 0.001$, ****$P < 0.0001$, ns: not significant.
